# A State Space Model of a Visuomotor Rotation Experiment

**DOI:** 10.1101/197483

**Authors:** Mireille E. Broucke

## Abstract

We present a new linear time-invariant (LTI) state space model to explain adaptation in human motor control. We focus on a visuomotor rotation experiment in which a human subject must rapidly move a cursor on a horizontal screen through a target disk. The hand itself is occluded from view, while the cursor is rotated relative to the hand angle by an amount of r degrees. Our model is based on the application of well-known techniques from control theory, in particular regulator theory. The model is simple, yet it reveals a plausible architecture for the high level computations underlying human motor control, including a representation of the internal model. It is a two state LTI model, where each state has a physical interpretation

## 1 Introduction

We present in this paper a new linear time-invariant (LTI) state space model to explain adaptation in human motor control. We focus on a visuomotor rotation experiment in which a human subject must rapidly move a cursor on a horizontal screen through a target disk. The hand itself is occluded from view, while the cursor is rotated relative to the hand angle by an amount of *r* degrees. There has been an outpouring of interest in such experiments on motor adaptation; see the bibliography where we include a sample of papers in the area.

Our model is based on the application of wellknown techniques from control theory, in particular regulator theory [12, 15, 64]. The model is simple, yet it reveals a plausible architecture for the high level computations underlying human motor control. Our model makes explicit how the internal model is updated [41, 26, 61]. It is a two state LTI model, where each state has a physical interpretation.

Our model is most closely related to the seminal work by Smith, Ghazizadeh, and Shadmehr [50]. They introduced a two state discrete time LTI model motivated by a viscous curl force field adaptation experiment (rather than the visuomotor rotation experiment considered here) in which one state corresponds to a slow mode and another state corresponds to a fast mode. Their two rate model reproduces a wide range of dynamic behaviors in motor adaptation. They assumed a generic structural form, resulting in an abstract model in which the states have no clear physical meaning, and the architecture of the computations remained unclear [34]. In contrast, our model gives physical significance to the fast and slow modes, and it provides architectural insight. Our findings put into question the hypothesis put forward by others that LTI models may be insufficient to capture all aspects of short term motor adaptation [65]. Like [50], our LTI model is able to recover savings with counterperturbation, savings with washout, reduced savings, anterograde interference, rapid unlearning, rapid downscaling, and spontaneous recovery. In addition, our model reveals how to reconcile error clamp behavior with the other dynamic behaviors of adaptation using only LTI models we recover new behaviors recently discovered using the error clamp paradigm [27]. We argue that, in general, linear models should be preferred over nonlinear models if they can recover the same behavior because they are simpler. Finally, while it is well accepted that internal models play a role in motor adapation [63], there is still debate on what is the exact form of the internal model. The internal model in our model takes the form of an *observer* [41].

Our model has a number of limitations. First, it only deals with highlevel motor behavior. As such, we do not consider the low, neuronal level, the exact signalling in the brain, or the precise role of the cerebellum. A second limitation is that our model is in continuous time, whereas the experiment itself evolves over discrete trials. This limitation is easily overcome: it is straightforward to generate an analogous discrete time equivalent of our continuous time model. Moreover, in control theory almost all results carry over seamlessly from the continuous domain to the discrete one, and vice versa. Thirdly, our model does not address all aspects of motor learning. In the terminology of [25], we deal with the “adaptation-only protocol”, and we consider adaptation for only one motor task at a time, each with its own associated internal model. Our model does not address how the brain assesses the relevance of errors [29] or how the degree of uncertainty about errors affects learning rates [5]. It does not deal with learning multiple tasks in parallel or switching between different internal models [34, 62]. Finally, it does not address the role of explicit strategies in implicit adaptation [39, 55].

This paper is organized as follows. First in Section 2 we describe in more detail the visuo motor rotation expriment and the eight dynamic behaviors that we focus on. In Section 3 we formulate, from a control-theoretic point of view, the problem of fast reach to a target in the presence of a disturbance. In Section 4 we present our solution to this problem in the form of a two state LTI model. This model is our main result. We give numerical simulations that show that our model is able to recover the first seven dynamic behaviors. In Section 4.1 we derive the error clamp model from the non-clamp model. We show it recovers the error clamp behavior reported in [27]. In Section 5 we explain the process by which we arrived at our model. This section is mathematically involved, and some background in control theory will be useful. The casual reader may skip Section 5 without any loss of understanding.

We use regulator theory to iteratively arrive at our solution. A first application of regulator theory in Section 5.2 yields a basic single state model that is already well known to be insufficient to capture all dynamic behaviors. In Section 5.3 we prove rigorously this model is inadequate, primarily to corroborate our theoretical approach with experimentally known facts. Next, in Section 5.4 we explore an alternative model inspired by the concept of an observer to obtain a twostate model that is capable of producing more behaviors. However, we show in Section 5.4 that this model still falls short of the requirement to reproduce all eight behaviors. A second application of regulator theory using an augmented open-loop model leads to the final model presented in Section 4.

## 2 Visuomotor Rotation Experiment

We consider an experiment to make a human rapidly reach for a small target disk on a planar surface. The subject is seated in front of a horizontal screen on which is displayed a disk representing the start position and a disk representing the target position. The instantaneous hand position is represented by a cursor on the screen. The hand itself is occluded from the subject’s view. The subject must move a mouse or stylus underneath the screen by observing the cursor on the screen. The task is to rapidly move the cursor from the starting position through the target disk. In the visuomotor rotation experiment, the placement of the cursor is rotated relative to the actual hand placement by an amount of *r* degrees. This rotation of the true hand position is thought to activate an error correction or socalled adaptation mechanism in the brain by presenting the subject with an evident and large error between the hand placement and the target disk. This mechanism of adaptation is explored through a number of variations of this basic experiment.

Visuomotor rotation experiments consist of sequences of blocks of trials of specific types in order to elicit certain dynamic behaviors. The types of blocks include:

1. **Baseline** (B). An initial block of trials when the subject is being familiarized with the aparatus and *r* = 0.
2. **Learning** (L). The first block of trials after the baseline block with *r*≠0.
3. **Washout** (W). A block of trials following a learning block with *r* = 0.
4. **Unlearning** (U). A block of trials following a learning block in which the rotation is set to -*r*.
5. **Relearning** (R). A second learning block. Typically, a washout or unlearning block is inserted between the first and second learning blocks.
6. **Downscaling** (D). A second learning block in which the rotation is set to a fraction of its value in the first learning block.
7. **Error Clamp** (E). There are at least two variants of the error clamp paradigm. In one variant, the cursor angle is pegged at the target disk angle (generally assumed to be zero) throughout the block. In the second variant, the cursor angle is pegged at the rotation value *r*.

A typical experiment proceeds in blocks of a prespecified order. For example, a BLUW experiment consists of a baseline block, a learning block, an unlearning block, and a washout block, in this order. The number of trials in each block can also be important. For example, a B_50_L_100_U_30_W_100_ experiment consists of 50 trials in the baseline block, 100 trials in the learning block, 30 trials in the unlearning block, and 100 trials in the washout block. When blocks of trials are sequenced in a particular order and with a particular number of trials in each block, then several phenomena emerge in experiments:

- *Savings* is a behavior in which learning is sped up in the second learning block relative to the first one. Two experiments in which savings can be exhibited are BLUR or BLWR.
- *Reduced savings* is a behavior in which savings is reduced by inserting a washout block of trials after the unlearning block. After the washout block, relearning does not proceed as rapidly as in the savings experiment. An experiment in which reduced savings may be exhibited is BLUWR.
- *Anterograde interference* is a behavior in which a previously learned task reduces the rate of subsequent learning of a different (and usually opposite) task. An experiment in which anterograde interference may be exhibited is BLU.
- *Rapid unlearning* is a behavior in which the rate of unlearning is faster than the rate of initial learning, if the number of trials in the learning block is small. An experiment in which rapid unlearning may be exhibited is a BLW experiment.
- *Rapid downscaling* is a behavior in which the rate of learning in a secondary learning block is faster when the rotation is set to a fraction of its value in the initial learning block. An experiment in which rapid downscaling may be exhibited is a B_50_L_50_D_50_ experiment with *r* = -30 in the learning block and *r* = −15 in the downscaling block.
- *Spontaneous recovery* is a behavior observed during the washout block of a BLUW experiment in which the hand angle partially “rebounds” to its value at the end of the learning block rather than converging monotonically to zero.
- *Error Clamp Behavior*. When the cursor angle is clamped at a fixed value, several interesting phenomema arise; these will be discussed in more detail in Section 4.1.

The goal of this paper is to explore the existence of a single LTI model that predicts all these behaviors. Our methodology is as follows. We first formulate a mathematical control problem that captures the problem of fast reach to a target disk. We then solve this problem using regulator theory, a body of results in control theory to design tracking controllers in the presence of unknown disturbances [64]. Inspired by this solution, we propose a hierarchy of models from simplest to most complex, until we arrive at a model that can explain the observed phenonema.

## 3 Problem Statement

We define several variables. Integer *n* is the number of reaches performed by the subject; *x*(*n*) is the angle (in degrees) of the final hand position at the nth reach relative to a reference line and measured at a predetermined radius from the start position; *r* is a constant rotation (in degrees) in the observed hand position; and

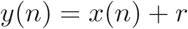

is the cursor angle and the measurement of the hand angle, as observed by the subject. Finally, we let 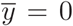 be the reference angle of the target disk. We assume that without a rotation, i.e. *r* = 0, the brain calibrates for a reach model of the form *x*(*n*) = 0; that is, the hand-eye coordination system is accurate and fast enough to produce a prelearned, nearperfect reach at each trial. Hence, mathematically speaking (under ideal conditions), there are no dynamics, no issues of stability, and the steadystate error is zero.

To stimulate an adaptation task, we introduce an unknown (to the subject) non-zero rotation so that the subject observes a cursor angle *y*(*n*) corresponding to a rotated hand position. The new model is given by:

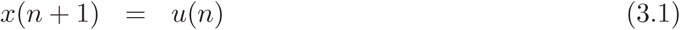

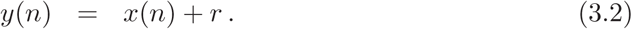

The model (3.1) states that *x*(*n* + 1), the reach at the (*n* + 1)-th attempt, is a function of information at the previous reach only. Therefore, we write the control input as *u*(*n*) (we could also write x(*n* + 1) = *u*(*n* + 1) with little difference of meaning). Because *u*(*n*), the *control input*, is a free variable that we choose, the model (3.1) places essentially no restrictions on the possible dynamics of the hand angle. Even the assumption that only information from the previous reach will be used is only enforced if the control input is a static measurement feedback (that is, an algebraic function of *y*(*n*)). If the control input is designed to be a dynamic feedback compensator, then the restriction is effectively removed. Regulator theory will inform us on what class of feedbacks is required.

In this paper we work with a continuous time model; a future report will deal with the discrete time case. Thus, instead of proceeding with the discrete model (3.1)-(3.2), we consider an equivalent continuous time model:

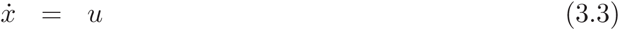

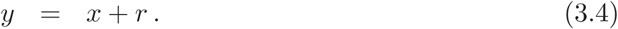

Here *x* ∈ ℝ is called the *state*, *u* ∈ ℝ is called the *control input*, and y is called the *output* or *measurement*. The meaning of (3.3) is that the brain has sufficient efficacy to assign the rate of change of the hand angle at will.

This continuous time model is to be regarded as an abstract model, but its variables correspond to the physical variables in the visuomotor experiment as follows: *x* is the angle of the hand, interpreted in a continuous time sense; *u* is a commanded rotational velocity of the hand; and *y* is the observed cursor angle. We define the *target error* to be

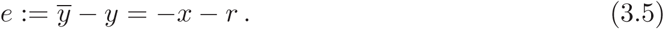

Researchers posit that by introducing an unknown rotation, the habitual hand-eye coordination system is derailed, and a new adaptation process is invoked. This process must satisfy certain requirements to be feasible: (i) it must be stable; (ii) it must generate a compensatory control strategy to asymptotically reach the target disk with small steady-state error. Additionally, we require that any proposed model will reproduce the behaviors informally described in Section 5.1. Thus, we are lead to the following control design problem.

**Problem 3.1.** Consider the system (3.3)-(3.4), where *r* is a constant, unknown rotation, and *y* is the measurement. Let 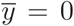 be the reference angle of the target disk. Find a measurement feedback control *u* = *f* (*y*) such that the closed-loop system is asymptotically stable and the target error satisfies

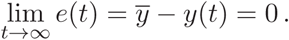

Moreover, the closed-loop system output *y* exhibits savings with counter perturbation, savings with washout, reduced savings, anterograde interference, rapid unlearning, rapid down-scaling, spontaneous recovery, and certain error clamp behavior. ⊳

## 4 Solution

In this section we present a solution to Problem 3.1 and we show via numerical simulations that the resulting model exhibits all the phenomena observed in the visuomotor experiment. At this stage we do not provide justifications on how this model was obtained. Our derivation of the model and several attempts that failed are deferred to Section 5.

We consider an error feedback of the form:

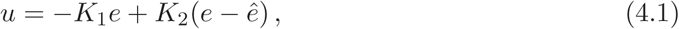

where *K*_1_, *K*_2_ ∈ ℝ are constants, and ê; is an estimate of the target error *e.* If we apply (4.1) to (3.3), then we get the closed-loop hand angle dynamics:

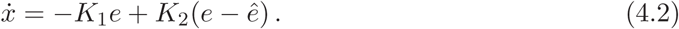

The estimate ê; is generated by an *observer* or *internal model* of the form:

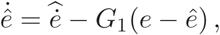

where *G*_1_ ∈ ℝ is a constant and 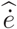 is an estimate of the derivative of the target error (which is not the same as the derivative of the target error estimate denoted by 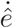). To obtain an estimate of the derivative of the target error, we recall that *e* = *-x - r* and *r* is a constant. Therefore, using (4.2) we have

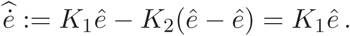

Using this formula, a reasonable estimate of ė is

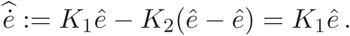

Using this estimate, we obtain the internal model of the target error:

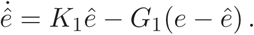

In summary, the proposed model is:

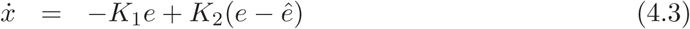

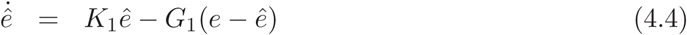

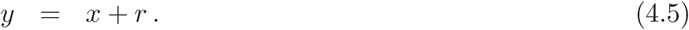

This LTI model consists of two states: the hand angle *x* and the estimate of the target error *ê*. The first equation models the dynamics of the hand angle over successive trials. If accurate, this model must be able to reproduce what is observed in experiments. The second equation comprises the internal model of the target error estimate, which is believed to arise out of activity in the brain. The term *G_1_(e-−ê)* guarantees that the target error estimate *ê* converges to the true target error *e*. The output as well as measurement of the model is the cursor angle *y*.

Due to the simplicity of this model and under the assumption that *r* is constant, it is possible to compute the solution explicitly. Define the *estimation error*

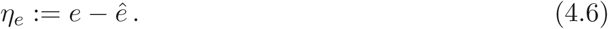

Then

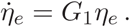

The solution of this differential equation is

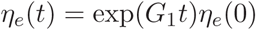

where *η*_*e*_(0) ∈ ℝ is the initial condition for *η*_*e*_. From this solution we get

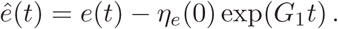

That is, the target error estimate converges exponentially to the true target error. Substituting the solution for *η*_*e*_(t) into (4.4) and using e = −*x* − *r*, we get

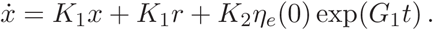

Assuming *r* is a constant and *K*_1_≠*G*_1_, the solution of this equation is:

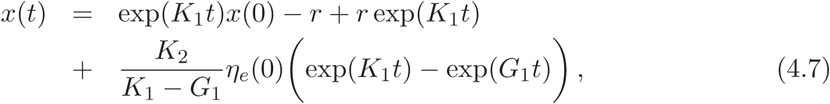

where *x*(0) is the initial condition for *x*. If we assume that the closed-loop system is stable,i.e. *K*_1_ < 0 and *G*_1_ < 0, then the steady-state solution is *x*_*ss*_ ≔ lim_*t*→∞_ *x*(*t*) = −*r* and *y*_*ss*_ = 0. Also, the steadystate error is *e*_*ss*_ = lim_*t*→∞_ −*x*(*t*)− *r* = 0. As such, the asymptotic behavior of this model is the one we expect. Figure 1 shows the behavior when *r* takes a sequence of three values: -30, 30, and 0 corresponding to an LUW experiment.

**Figure 1:**
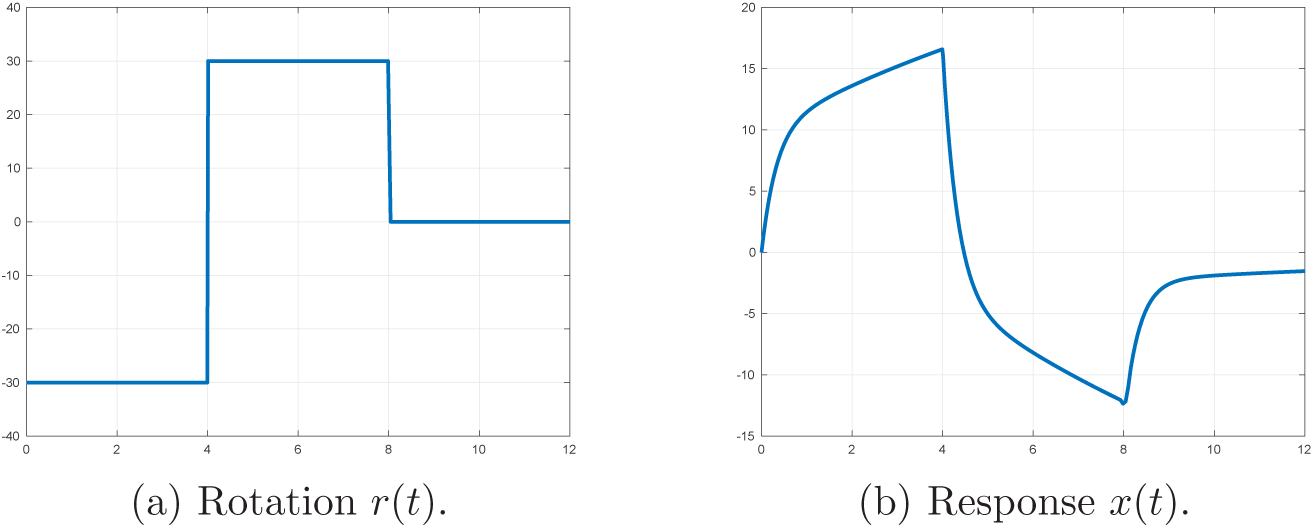
Behavior of (4.3)-(4.5) for an LUW experiment.

By observing the form of the solution, we see that, as with the model in [50], our model is a two rate model. In simulation we will see that the slow mode corresponds to the hand angle dynamics while the fast mode corresponds to the rate at which the error estimate converges to the error. More precisely, the rate of the slow mode is determined by the parameter *K*_1_ while the fast mode is determined by the parameter *G*_1_. An interesting idea is that the two rate behavior appears to arise from the interaction between a slow process in the observable world and a fast process retained in the brain.

Figures 2-8 show results from numerical simulations demonstrating that our model generates the first seven dynamic behaviors. For all simulations the model parameters are fixed as follows: *K*_1_ = −1/10, *K*_2_ = 1, and *G*_1_ = −2.

**Figure 2:**
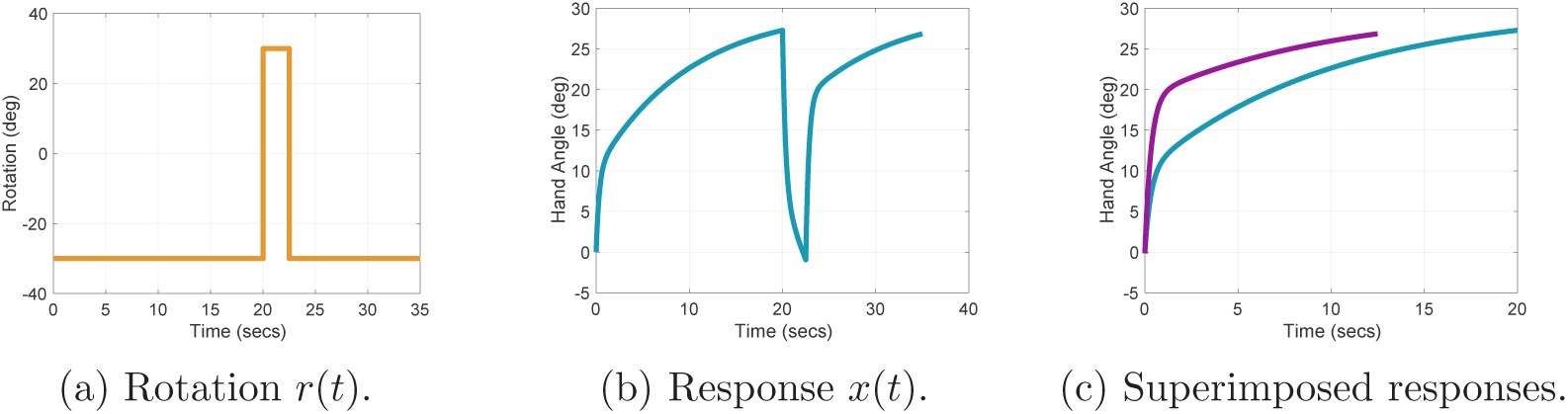
Savings with CP exhibited by (4.3)-(4.5) for an LUR experiment. The left figure shows the rotation as a function of time: *r(t)* = −30 during the learning block, *r(t) =* +30 during the unlearning block, and *r(t) = -*30 during the relearning block. The center figure shows the hand angle where it is observed that the rate that *x*(*t*) approaches its steady-state value *x*_*ss*_ = 30 is faster in the relearning block than in the learning block. This fact is verified in the right figure where we plot *x*(*t*) during the learning block superimposed with a shifted version of *x*(*t*) during the relearning block. Precisely, *x*(*t*) over the time interval t ∈ [0, 20) is shown in blue, and *x*(*t* +*t*_3_) over the time interval t ∈ [0, 13) is shown in purple. The time *t*_3_ is the third time when *x*(*t*) equals 0. We can see that the purple curve is larger than the blue curve, corresponding to faster learning in the relearning block.

**Figure 3:**
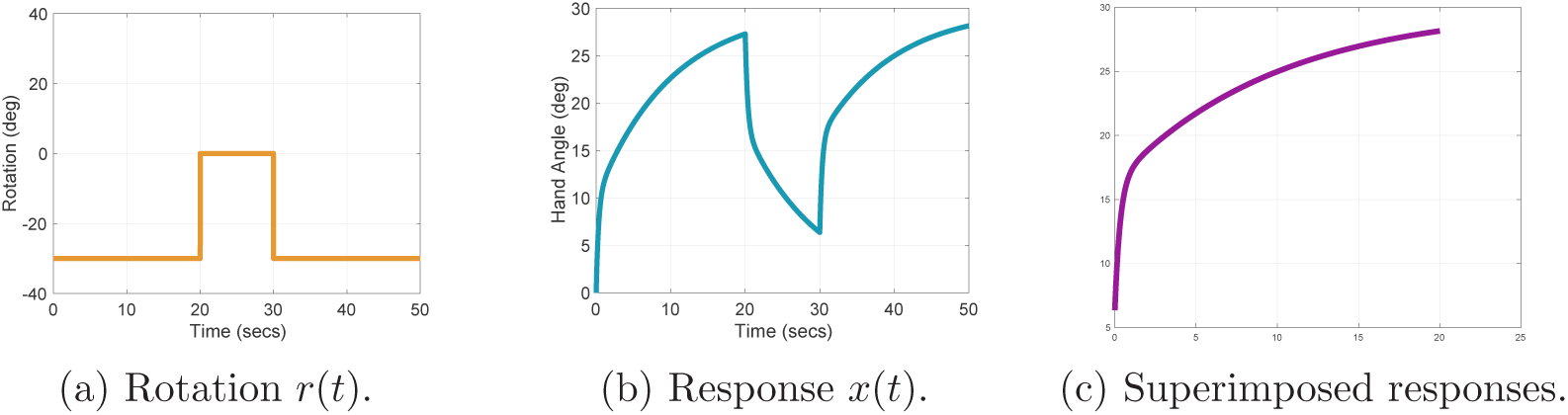
Savings with WO exhibited by (4.3)-(4.5) for an LWR experiment. The left figure shows the rotation as a function of time: *r*(*t*) = −30 during the learning block, *r*(*t*) = 0 during the washout block, and *r*(*t*) = −30 during the relearning block. The center figure shows the hand angle where it is observed that the rate that *x*(*t*) approaches its steady-state value *x*_*ss*_ = 30 is faster in the relearning block than in the learning block. In the right figure *x*(*t* + *t*_1_) over the time interval of the learning block is shown in blue, and *x*(*t* + *t*_2_) over the time interval of the relearning block is shown in purple. The time *t*_1_ near the beginning of the learning block and the time *t*_2_ near the beginning of the relearning block are selected such that *x*(*t*_1_) = *x*(*t*_2_). We can see that the purple curve is larger than the blue curve, corresponding to faster learning in the relearning block.

**Figure 4:**
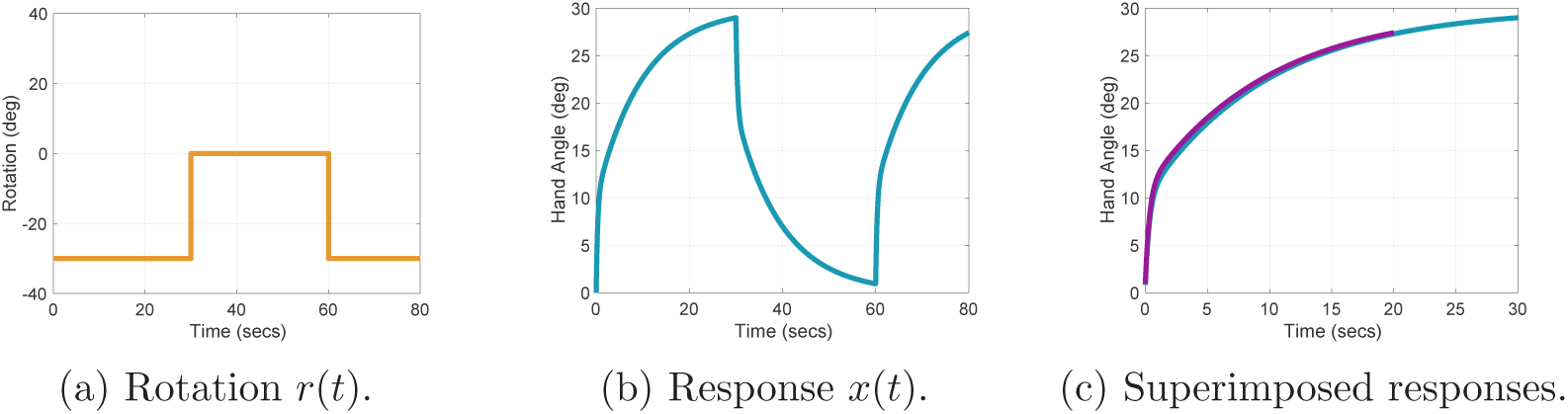
Reduced savings exhibited by (4.3)-(4.5) for an LWR experiment. The left figure shows the rotation as a function of time: *r*(*t*) = −30 during the learning block, *r*(*t*) = 0 during the washout block, and *r*(*t*) = −30 during the relearning block. The center figure shows the hand angle where it is observed that the rate that *x*(*t*) approaches its steady-state value *x*_*ss*_ = 30 is roughly the same as in the relearning block. In the right figure *x*(*t* + *t*_1_) over the time interval of the learning block is shown in blue, and *x*(*t* + *t*_2_) over the time interval of the relearning block is shown in purple. The time *t*_1_ near the beginning of the learning block and the time *t*_2_ near the beginning of the relearning block are selected such that *x*(*t*_1_) = *x*(*t*_2_). We can see that the purple curve is almost identical to the blue curve, so savings has been reduced.

**Figure 5:**
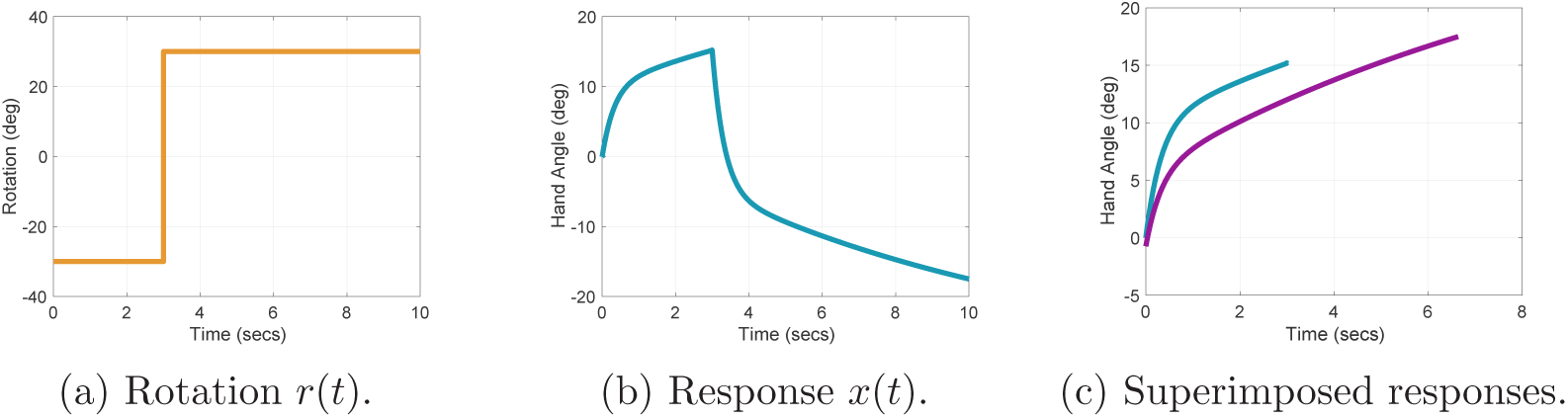
Anterograde interference exhibited by (4.3)-(4.5) for an LU experiment. The left figure shows the rotation as a function of time: *r*(*t*) = −30 during the learning block, and *r*(*t*) = 30 during the unlearning block. The center figure shows the hand angle where it is observed that the rate that *x*(*t*) approaches its steady-state value *x*_*ss*_ = −30 in the unlearning block is slower than the rate that it approaches the steadystate value *x*_*ss*_ = 30 in the learning block. In the right figure *x*(*t*) over the time interval of the learning block is shown in blue, and −*x*(*t* + *t*_2_) over the time interval of the unlearning block is shown in purple. The time *t*_2_ at the beginning of the unlearning block is selected such that *x*(0) = *x*(*t*_2_) = 0. We can see that the blue curve is larger than the block curve, indicating that the learning rate is reduced in the unlearning block.

**Figure 6:**
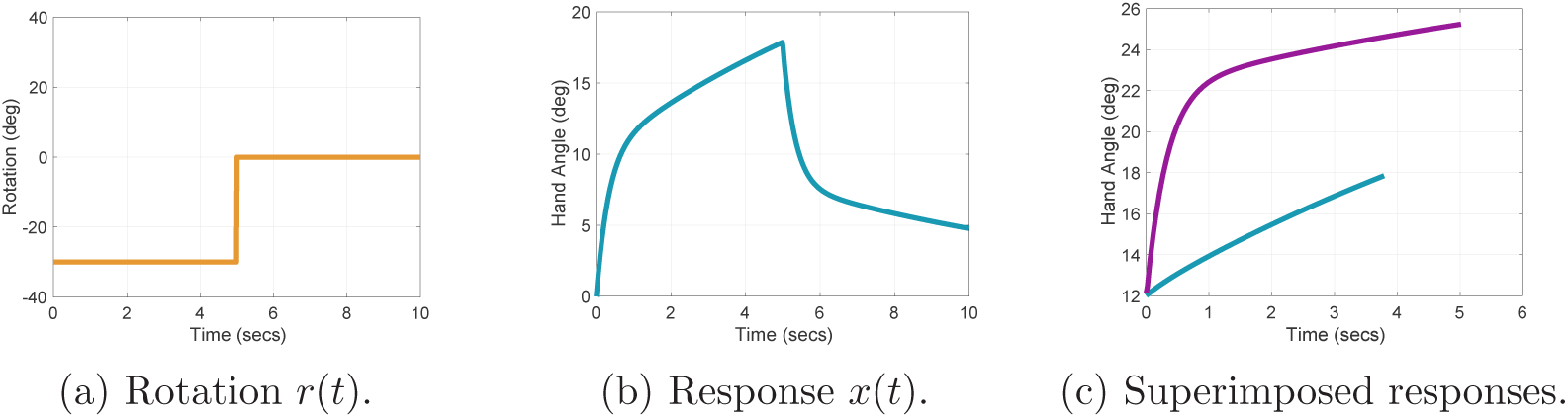
Rapid unlearning exhibited by (4.3)-(4.5) for an LW experiment. The left figure shows the rotation as a function of time: *r*(*t*) = −30 during the learning block, and *r*(*t*) = 30 during the washout block. The center figure shows the hand angle where it is observed that the rate that *x*(*t*) approaches its steady-state value *x*_*ss*_ = 30 in the learning block is slower than the rate that it returns to the baseline value of 0 in the washout block. In the right figure *x*(*t*+*t*_1_) over the time interval of the learning block is shown in blue, and 30−*x*(*t*+*t*_2_) over the time interval of the unlearning block is shown in purple. The time *t*_1_ is selected so that *x*(*t*_1_) = 30 − *x*(*t*_2_), while *t*_2_ is the start time of the washout block. We can see that the purple curve is greater than the blue curve at each time point, indicating that the rate of washout is greater than the rate of learning.

**Figure 7:**
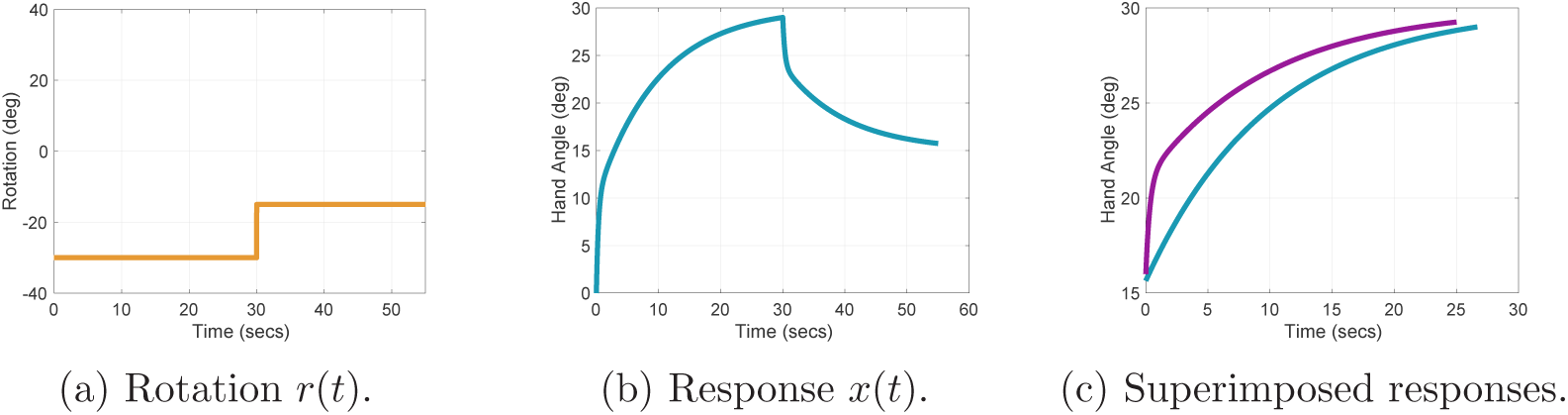
Rapid downscaling exhibited by (4.3)-(4.5) for an LD experiment. The left figure shows the rotation as a function of time: *r*(*t*) = −30 during the learning block, and *r*(*t*) = −15 during the downscaling block. The center figure shows the hand angle where it is observed that the rate that *x*(*t*) approaches its steady-state value *x*_*ss*_ = 30 in the learning block is slower than the rate that it returns to the downscaled value of 15. In the right figure *x*(*t*+*t*_1_) over the time interval of the learning block is shown in blue, and 45−*x*(*t*+*t*_2_) over the time interval of the unlearning block is shown in purple. The time *t*_1_ is selected so that *x*(*t*_1_) = 45 − *x*(*t*_2_), while *t*_2_ is the start time of the downscaling block. We can see that the purple curve is greater than the blue curve at each time point, indicating that the rate of downscaling is greater than the rate of learning.

**Figure 8:**
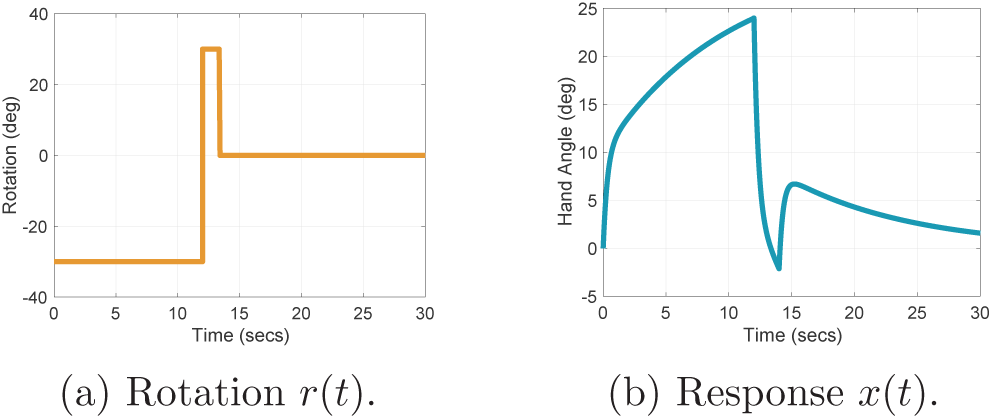
Spontaneous recovery exhibited by (4.3)-(4.5) for an LUW experiment. The left figure shows the rotation as a function of time: *r*(*t*) = −30 during the learning block, *r*(*t*) = 30 during the unlearning block, and *r*(*t*) = 0 during the washout block. The right figure shows the hand angle *x*(*t*). We observe that during the washout block corresponding to *t* ∈ [14, 30], the hand angle rebounds to a value greater than zero, even though the steady state value for the washout block is *x*_*ss*_ = 0.

### 4.1 Error Clamp Model

In the error clamp paradigm the cursor angle is pegged at *r* [27]. The measurement becomes

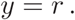

We examine the effect of pegging the measurement at a constant value on the internal model. Because there is no perceived change in the target error, the estimate of the target error derivative 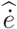 is zero. The new internal model equation is:

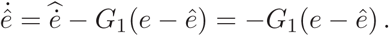

The effect of pegging the measurement at a constant value on the hand angle dynamics is more subtle. First, because *e* = −*x* − *r* in non-clamp conditions, (4.3) can be re-written as:

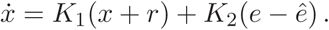

During the error clamp, information about the hand angle is lost. This is reflected through the fact that we replace *r* by 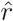, a constant parameter with no apparent relationship to *r*. In summary, the error clamp model is:

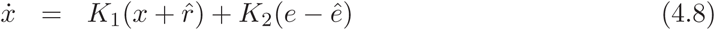

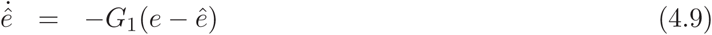

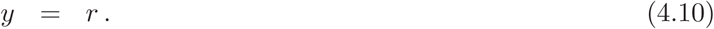

Simulation results with the error clamp model are shown in Figure fig:errorclamp. The parameter values are: *K*_1_ = −1/10, *K*_2_ = 1, *G*_1_ = −2 and 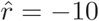. Clearly more research is needed to understand the underlying mechanism for the loss of state information in the error clamp paradigm.

## 5 Derivation

In this section we show how our model was derived. First we present formal mathematical definitions of the dynamic behaviors so that it is possible later to prove rigorously that certain simpler models do not reproduce these behaviors. Second we apply regulator theory to solve Problem 3.1. Background on regulator theory is provided in the Appendix. Next, we give two intermediate solutions to the problem and show that they do not reproduce all behaviors. These findings ultimately lead us to the final model of Section 4.

**Notation.** We use the following notation. The spectrum or set of eigenvalues of a matrix *A* ∈ ℝ^*n*×*n*^ is denoted σ(*A*). The set of complex numbers is denoted ℂ, and ℂ^−^ ≔ {λ ∈ ℂ │ ℜ(λ) < 0} denotes the open left half complex plane. Given a function w : ℝ → ℝ^*q*^ and a constant vector *r* ∈ ℝ^*q*^, we write *w*(*t)* ≡ *r* if w(t) = *r* for all *t* ∈ ℝ.

### 5.1 Dynamic Behaviors

In this section we formalize the notions of savings, reduced savings, anterograde interference, rapid unlearning, rapid downscaling, and spontaneous recovery.

Consider a general LTI model with a single output given by

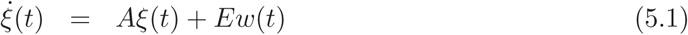

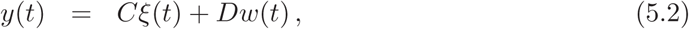

where *ξ*(*t*) ∈ ℝ^*n*^ is the state, *w*(*t*) ∈ ℝ^*q*^ is a disturbance, and *y*(*t*) ∈ ℝ is the measurement or output. Suppose that the system is asymptotically stable, i.e. σ(*A*) ⊂ ℂ^−^. In the following definitions, let *r* ∈ ℝ^*q*^, *y*_0_ ∈ ℝ, and *t*_0_ ≥ 0 be constants. Let *y*_*ss*_ be the steadystate value of *y* with initial value *y*(*t*_0_) = *y*_0_ when *w*(*t*) ≡ *r*. Also we assume that −*y*_*ss*_ is the steady-state value of *y* with initial value *y*_0_ when *w*(*t*) ≡ −*r*. Observe that if *w* = 0, then the steadystate value of *y* is *y*_*ss*_ = 0 because the system is stable.

**Definition 5.1.** Consider the stable LTI system (5.1)-(5.2), and let *r* ∈ ℝ^*q*^, *y*_0_ ∈ ℝ, *t*_0_ ≥ 0, and *y*_*ss*_ ∈ ℝ be as above. We define the *rise time*, denoted *T*_*r*_, to be the time duration for *y*(*t*) to reach 90% of its steady-state value *y*_*ss*_, starting from an initial value *y*(*t*_0_) = *y*_0_. ⊳

**Definition 5.2.** Consider the stable LTI system (5.1)-(5.2), and let *r* ∈ ℝ^*q*^, *y*_0_ ∈ ℝ, *t*_0_ ≥ 0, and *y*_*ss*_ ∈ ℝ be as above. Let *y*_*max*_ = max_*t*≥*t*_0 *y*(*t*). The *overshoot* of *y*(*t*) is defined to be OS ≔ │*y*_*max*_│ - │*y*_*ss*_│. ⊳

Savings corresponds to a reduction in the rise time in the subsequent evolution of *y*(*t*) starting from an initial value *y*_0_.

**Definition 5.3** (Savings). Consider the stable LTI system (5.1)-(5.2), and let *r* ∈ ℝ^*q*^, *y*_0_ ∈ ℝ, *t*_0_ ≥ 0, and *y*_*ss*_ ∈ ℝ be as above. Suppose we are given times t_3_ ≥ t_2_ > t_1_ > t_0_ such that: *w*(*t*) = *r* for *t* ∈ [*t*_0_, *t*_1_) ∪ [*t*_2_, ∞), and *y*(*t*_3_) = *y*(*t*_0_) = *y*_0_. Let *T*_*r*0_ and *T*_*r*3_ be the rise times starting at *t*_0_ and *t*_3_, respectively. We say (5.1)-(5.2) exhibits *savings* if

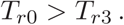

Additionally, if *w*(*t*) = −*r* for *t* ∈ [*t*_1_, *t*_2_), then we say (5.1)-(5.2) exhibits *savings with counter perturbation (CP)*. If *w*(*t*) = 0 for *t* ∈ [*t*_1_, *t*_2_), then we say (5.1)-(5.2) exhibits *savings with washout (WO)*. ⊳

*Remark* 5.1. We observe that Definition 5.3 makes no explicit statements about the rate of increase or decrease of *y*(*t*). The rates of exponentials generated by the LTI model (5.1)-(5.2) are fixed by the eigenvalues of *A*. What varies between the first and second learning blocks are the initial conditions *ξ*(*t*_0_) and *ξ*(*t*_2_) which determine the linear combination of exponentials appearing in *y*(*t*) in each block. This linear combination then determines the change in rise time, in turn leading to the possibility of savings. Second, notice that Definition 5.3 is ambiguous about the choice of *y*_0_. Typically, we take *t*_0_ = 0, so *y*_0_ = *y*(0), and *t*_3_ is the first time after *t*_2_ when *y*(*t*_3_) = *y*_0_. Finally, we note that it need not be the case that *y*(*t*) has already reached 90% of its steadystate value *y*_*ss*_ at time *t*_1_ when the first learning block ends. Rise time can still be computed by extending forward the given curve. Rise time is used here primarily because it is a standard metric in control theory to quantify speed of response to step inputs (in this case the step input is the disturbance *w*). Other metrics can also be used.

Reduced savings occurs when a block of washout trials is inserted between two blocks of learning trials under the same value of *w*.

**Definition 5.4** (Reduced Savings). Consider the stable LTI system (5.1)-(5.2), and let *r* ∈ ℝ^*q*^, *y*_0_ ∈ ℝ, *t*_0_ ≥ 0, and *y*_*ss*_ ∈ ℝ be as above. Also suppose we are given a duration *T*_*wo*_ > 0 and times *t*_2_ > *t*_*wo*_ + *T*_*wo*_ > *t*_*wo*_ > *t*_1_ > *t*_0_ such that: *w*(*t*) = *r* for *t* ∈ [*t*_0_, *t*_1_) ∪ [*t*_*wo*_ + *T*_*wo*_, ∞),

*w*(t) = 0 for *t* ∈ [*t*_*wo*_, *t*_*wo*_ + *T*_*wo*_), and *y*(*t*_2_) = *y*(*t*_0_) = *y*_0_. Let *T*_*r*0_ and *T*_*r*2_ be the rise times starting at *t*_0_ and *t*_2_, respectively. We say (5.1)-(5.2) exhibits *reduced savings* if *T*_*r*0_ ≥ *T*_*r*2_ and

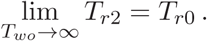

**Definition 5.5** (Anterograde Interference). Consider the stable LTI system (5.1)-(5.2), and let *r* ∈ ℝ^*q*^ and *y*_*ss*_ ∈ ℝ be as above. Also suppose we are given times *t*_2_ > *t*_1_ > *t*_0_ such that: *w*(*t*) = *r* for t ∈ [*t*_0_, *t*_1_), *w*(*t*) = −*r* for *t* ∈ [*t*_1_, ∞), and *y*(*t*_2_) = *y*(*t*_0_) = *y*_0_. Let *T*_*r*0_ and *T*_*r*2_ be the rise times starting at *t*_0_ and *t*_2_, respectively. We say (5.1)-(5.2) exhibits *anterograde interference* if

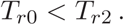

Moreover *T*_*r*2_ increases as the duration of the first learning block *t*_1_ increases. ⊳

*Remark* 5.2. Other definitions of anterograde interference can be considered. For example, suppose *t*_0_ = 0 and *y*(0) = 0. Let *T*_*r*0_ be the rise time of *y*(*t*) starting at *t* = 0, as above. Then consider

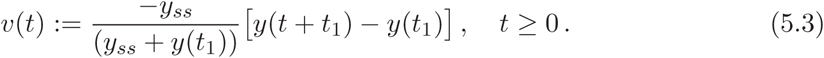

Let *T*_*rv*_ be the rise time of *v*(*t*) starting at *t* = 0. We say (5.1)-(5.2) exhibits *anterograde interference* if

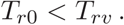

The signal *v*(*t*) in this definition corresponds to shifting *y*(*t*) so that *v*(0) = 0 and then rescaling so that *v*_*ss*_ = lim_*t*→∞_ *v*(*t*) = *y*_*ss*_ (recall by assumption lim_*t*→∞_ *y*(*t* + *t*_1_) = −*y*_*ss*_. This shifting and rescaling allows for a fair comparison with the rise time of *y*(*t*) in the first learning block by matching its initial condition *y*_0_ = 0 and its steady-state value *y*_*ss*_. However, care must be taken in applying this definition. For scalar LTI systems (when *n* = 1) the two definitions of anterograde interference will give the same result, but for general dynamical systems, the two definitions are very different. As an example, suppose the output has the form:

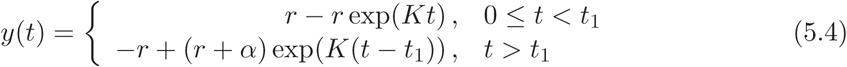

where *K* < 0 and α = *r* − *r* exp(*Kt*_1_). See Figure 10. Such an output could be generated by a scalar LTI system with two blocks of trials with different values of the disturbance *w*. Since the system generating this response is scalar, we expect that the two notions of anterograde interference will coincide. Particularly, we show neither notion of anterograde interference is exhibited.

**Figure 9:**
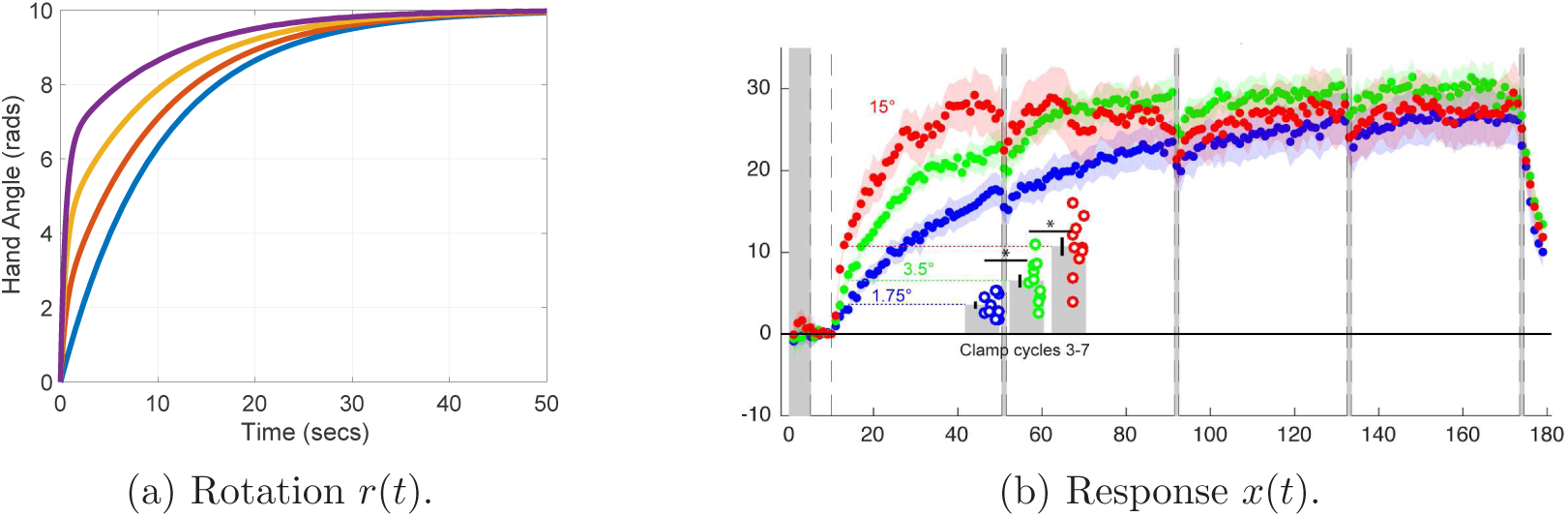
Error clamp behavior exhibited by (4.8)-(4.10). The left figure shows the hand angle when the error is clamped at 2, 8, and 15 degrees, respectively. We observe that the rate of adaptation increases with the magnitude of the clamped error, but the asymptotic value of *x* is invariant. The right figure is Figure 2 in [27] showing the hand angle as a function of trials for clamped errors 1.75, 3.5, and 15 degrees.

**Figure 10:**
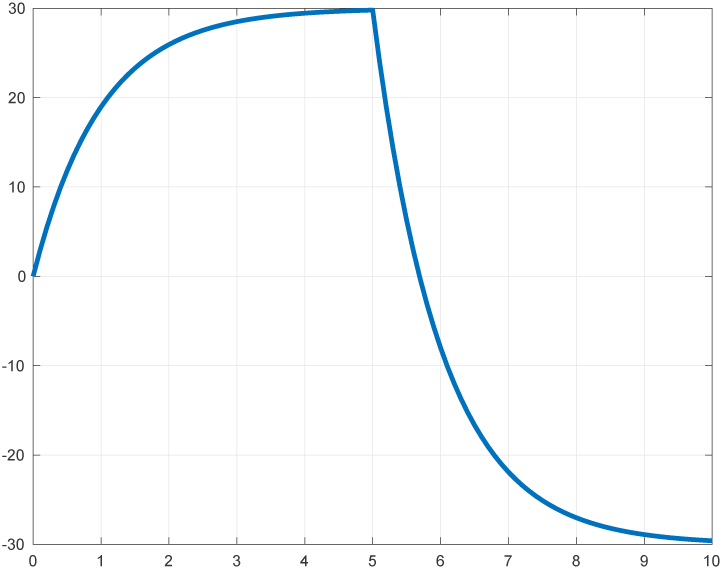
Response *x*(*t*) of (5.6)-(5.7) with *K* = −1 for an LUW experiment.

Regarding Definition 5.5, we have that *t*_0_ = 0 and *y*_0_ = *y*(*t*_0_) = 0. Then we must compare the rise time of *y*(*t*) on the time interval [0, *t*_1_) with the rise time of *v*(*t*) = −*y*(*t* + *t*_2_) on the time interval [*t*_2_, ∞), where *t*_2_ > *t*_1_ is such that *y*(*t*_2_) = *y*(*t*_0_) = *y*_0_. We have that

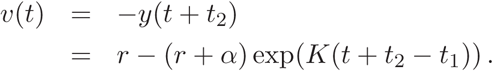

However, by assumption *y*(*t*_2_) = 0 = −*r*+(*r*+α) exp(*K*(*t*_2_ −*t*_1_)), so (*r*+α) exp(*K*(*t*_2_ −*t*_1_)) =*r*. Therefore,

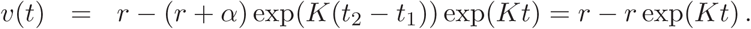

Since *v*(*t*) = *y*(*t*) for *t* ∈ [0, *t*_1_), this notion of anterograde interference can not occur.

Now consider the second notion. We have that *y*_*ss*_ = *r* and *y*(*t*_1_) = α. Substituting into (5.3), we obtain

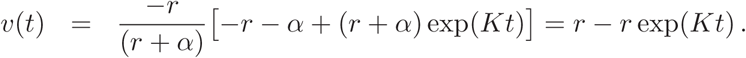

Again *v*(*t*) = *y*(*t*) for *t* ∈ [0, *t*_1_), so this notion of anterograde interference also can not occur.

**Definition 5.6** (Rapid Unlearning). Consider the stable LTI system (5.1)-(5.2), and let r ∈ ℝ^*q*^, *y*_0_ ∈ ℝ, *t*_0_ ≥ 0, and *y*_*ss*_ ∈ ℝ be as above. Suppose we are given times *t*_2_ > *t*_1_ > *t*_0_ such that: *w*(*t*) = *r* for t ∈ [*t*_0_, *t*_2_), *w*(*t*) = 0 for *t* ∈ [*t*_2_, ∞), and *y*(*t*_1_) = *y*_*ss*_ − *y*(*t*_2_). Let *T*_*r*1_ and *T*_*r*2_ be the rise times starting at *t*_1_ and *t*_2_, respectively. We say (5.1)-(5.2) exhibits *rapid unlearning* if

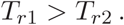

Moreover *T*_*r*2_ decreases as the duration of the first learning block *t*_2_ decreases. ⊳

**Definition 5.7** (Rapid Downscaling). Consider the stable LTI system (5.1)-(5.2), and let *r* ∈ ℝ^*q*^, *y*_0_ ∈ ℝ, *t*_0_ ≥ 0, and *y*_*ss*_ ∈ ℝ be as above. Suppose we are given α ∈ (0, 1) and times *t*_2_ > *t*_1_ > *t*_0_ such that: *w*(*t*) = *r* for *t* ∈ [*t*_0_, *t*_2_), *w*(*t*) = αr for *t* ∈ [*t*_2_, ∞), and *y*(*t*_1_) = (1 + α)*y*_*ss*_ − *y*(*t*_2_). Also, we assume that the steadystate value of *y*(*t*) for *t* ≥ *t*_2_ is α*y*_*ss*_, and *y*(*t*_2_) > α*y*_*ss*_. Let *T*_*r*1_ and *T*_*r*2_ be the rise times starting at *t*_1_ and *t*_2_, respectively. We say (5.1)-(5.2) exhibits *rapid downscaling* if

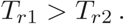

Moreover *T*_*r*2_ decreases as the duration of the first learning block *t*_2_ decreases. ⊳

*Remark* 5.3. The justification for the expression *y*(*t*_1_) = (1 + α)*y*_*ss*_ − *y*(*t*_2_) in the previous definition is as follows. To make a fair comparison between the rise times for the learning block and the downscaling block, the output *y*(*t*) must vary over the same range of values. If the initial time for the measurement of rise time in the learning block is selected to be *t*_1_, then the total variation of *y*(*t*) from this time is *y*_*ss*_ − *y*(*t*_1_). Similarly, for the downscaling block, the total variation of *y*(*t*) is *y*(*t*_2_) - α*y*_*ss*_. Equating these two expressions and solving for *y*(*t*_1_), we obtain the expression above.

**Definition 5.8** (Spontaneous Recovery). Consider the stable LTI system (5.1)-(5.2), and let *r* ∈ ℝ^*q*^, *y*_0_ ∈ ℝ, *t*_0_ ≥ 0, and *y*_*ss*_ ∈ ℝ be as above. Suppose there exists a time *t*_1_ > *t*_0_ such that *y*(*t*_1_) = *y*(*t*_0_) = *y*_0_. We say (5.1)-(5.2) exhibits *spontaneous recovery* if the overshoot starting from *y*(*t*_1_) = *y*_0_ satisfies:

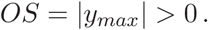

⊳

### 5.2 Regulator Theory I

In this section we apply regulator theory to solve Problem 3.1. It is helpful at this point to review the results in the Appendix, particularly Section A.1 on observer-based regulator design.

Comparing our model (3.3)-(3.4) with the general model (A.1), we have *A* = 0, *B* = 1, *E* = 0, *C* = 1, *D* = 1, and w = *r*. The *exosystem* provides a dynamic model for the constant rotation *r*. It has the form

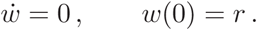

This model characterizes that it is known that the disturbance is constant; however, we do not know the initial condition *w*(0) = *r*. Comparing to the general exosystem model (A.2), we see that *S* = 0. Observe that σ(S) = {0} ⊂ ℂ^+^. Now we must verify conditions (N1) and (N2) to apply Theorem A.1. The pair (*A*, *B*) = (0, 1) is trivially controllable and therefore stabilizable. Similarly, the pair (*C*, *A*) = (1, 0) is trivially observable and therefore detectable. Next we verify (N3). We have

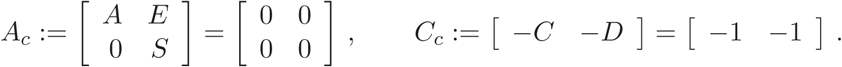

The observability matrix associated with the pair (*C*_*c*_, *A*_*c*_) is:

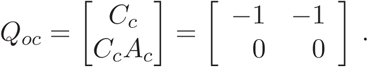

This matrix is not full rank so the pair (*C*_*c*_, *A*_*c*_) is not observable. It is also easy to check that (*C*_*c*_, *A*_*c*_) is not detectable since 0 ∈ ℂ^+^ is an unobservable eigenvalue. Now we apply the procedure in Lemma A.2 to prune the exosystem to remove redundant information. Then we can proceed with the regulator design for the pruned system. To find the observable part, note that the unobservable subspace is spanned by the single vector (1, −1). We select a first basis vector (1, 0) in order to retain the plant dynamics. Then a coordinate transformation that yields the observable and unobservable subsystems is 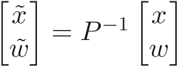, where

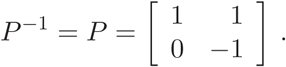

This yields a state for the observable subsystem to be 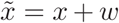. Since *w* is assumed to be a constant, we get 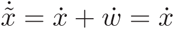. From this observation we obtain the reduced system

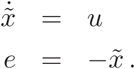

We observe that the exosystem has been completely eliminated. According to Remark A.1, the output regulation problem reduces to a problem of output stabilization. Namely, we must find an error feedback *u = f (e)* such that the closed-loop system is asymptotically stable. An immediate solution to this problem is to select

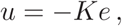

where *K < 0*. We analyze this solution in the next section.

### 5.3 Error Feedback

Based on the discussion in the previous section, the first model we consider uses *error feedback* of the form

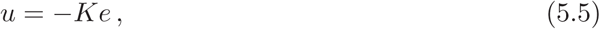

where *K* ∈ ℝ is a constant called the *loop gain*. The formula (5.5) makes the assumption that the target error is exactly the same error signal used in the feedback law. In reality, the subject likely generates an estimate of e or uses a noisy variant of e. Nevertheless, this first order model accounts for the dominant effect of using a signal that roughly approximates or is proportional to the target errror.

Applying (5.5) to (3.3) and using (3.5), the closedloop system is:

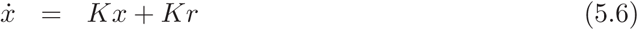

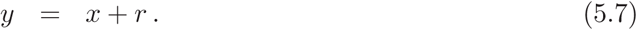

The solution of this equation is:

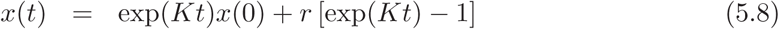

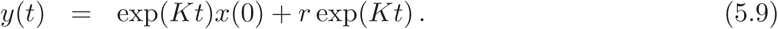

where *x(0)* is the initial condition. If we assume that the closedloop system is stable, i.e. *K* < 0, then the steadystate solution is *x*_*ss*_ *≔* lim_*t*→∞_ *x(t) = -r* and *y*_*ss*_ = 0. Also, the *steadystate error* is:

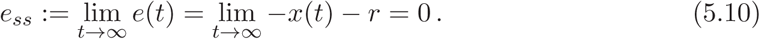

We can see that the asymptotic behavior of this model is the one we expect. Figure 11 shows the behavior when *r* takes a sequence of three values: −30, 30, and 0 corresponding to an LUW experiment (the initial conditions have been assigned to emulate a baseline block occurring before the learning block).

**Figure 11:**
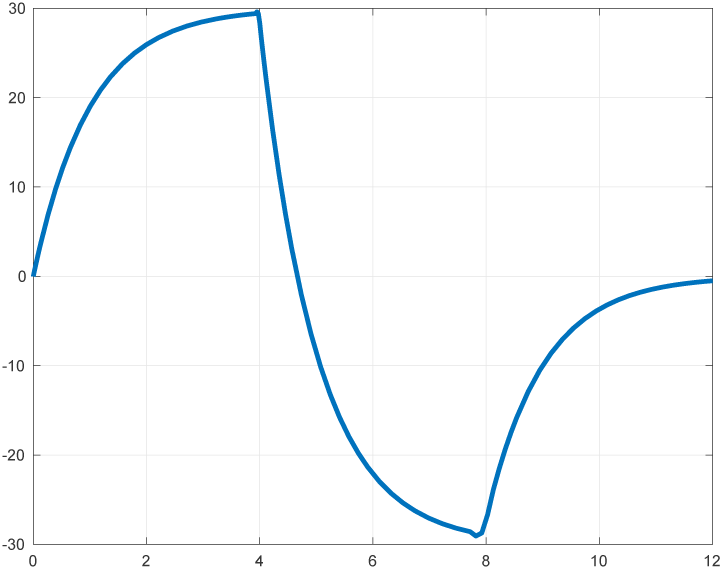
Response *x*(*t*) of (5.6)-(5.7) with *K* = −1 for an LUW experiment.

It is well-known that this error feedback model does not account for all the behaviors observed in the visuomotor rotation experiment. Here we focus on one behavior, savings, which can not be generated by this model.

**Lemma 5.1**. *The stable closed-loop system* (5.6)-(5.7) *does not exhibit savings.*

*Proof.* If we compare the model (5.1)-(5.2) with (5.6)-(5.7), we observe that *n* = 1, *q* = 1, *ξ* = *y* = *x*, *A* = *E* = *K*, *C* = 1, *D* = 0, and *w* = *r*. Suppose we are given *r* ∈ ℝ. Without loss of generality, suppose *x*(*t*_0_) = *x*_0_ = 0. Suppose there exist t_1_, *T*_*r*0_, *T*_*r*1_ > 0 such that *x*(*t*_0_) = *x*(*t*_1_) = 0 and *x*(*t*_0_ + *T*_*r*0_) = *x*(*t*_1_ + *T*_*r*1_) = 0.9*x*_*ss*_. Then applying (5.8) twice we have

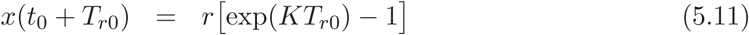

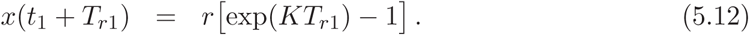

Subtracting (5.12) from (5.11) and dividing by *r*, we obtain exp(*KT*_*r*0_) = exp(*KT*_*r*1_),so│*KT*_*r*0_│=│*KT*_*r*1_│. That is, *T*_*r*0_ = *T*_*r*1_, so there is no savings. □

Figure 12 confirms the prediction of Lemma 5.1. Figure 12(a) shows the response when *r* takes a sequence of values −30, 30, and −30 corresponding to an LUR experiment. In Figure 12(b) the two responses for *r* = −30 are superimposed with both starting from the initial value of *x* = 0. We can see that the rate of response in the second block (shown in blue) is the same as the rate of response of the first block (show in red), so there is no savings.

**Figure 12:**
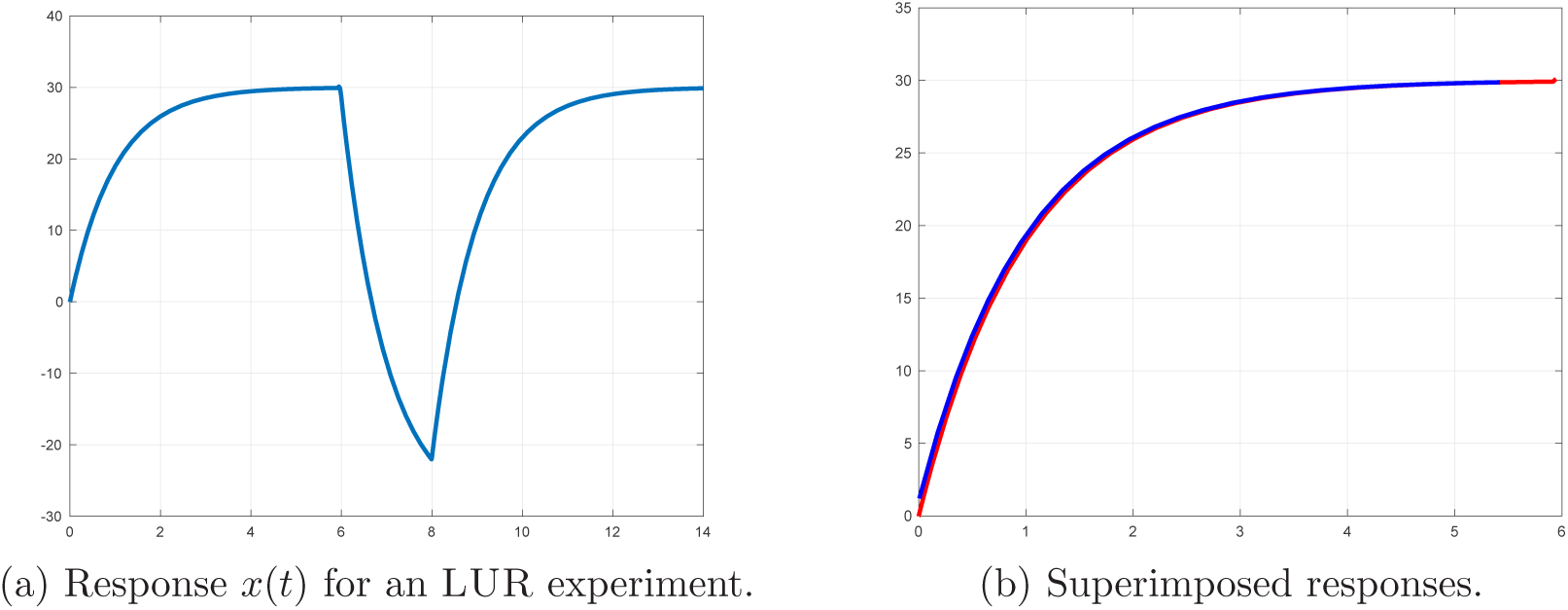
Savings experiment for (5.6)-(5.7).

In additional issue with the error feedback model is that it is not robust to plant perturbations. Suppose that the true hand dynamics have the form

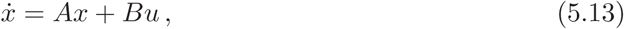

where *A*, *B* ∈ ℝ are unknown parameters. Suppose we again apply the error feedback *u* = −*Ke*, and we assume that *A* is sufficiently close to 0 and *B* is sufficiently close to 1

such that σ(*A* + *BK*) ⊂ ℂ^−^. Since *r* is constant, 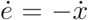. Then substituting *u* = −*Ke* in (5.13), we have

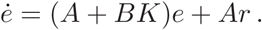

The solution of this differential equation is

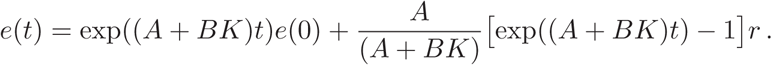

Because σ(*A* + *BK*) ⊂ ℂ^−^, we can compute the steadystate error:

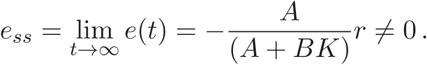

In sum, the error feedback controller is not robust to plant uncertainty because it is not capable to drive the target error to zero without exact knowledge of the plant parameters.

From the foregoing analysis we conclude the single state model is not adequate to capture the behavior in the visuomotor experiment. Error feedback is likely a component of a more sophisticated model.

### 5.4 Internal Model

In the second model we attempt to correct the shortcomings of the error feedback model. To that end, we augment the first order model by an internal model of the target error. We consider an error feedback of the form:

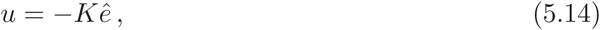

where *K* ∈ ℝ is the same as before, whilê is an estimate of the target error rather than the target error itself. The estimat ê is generated by an *observer* or *internal model*. The observer equation is given by

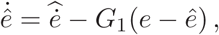

where *G*_1_ ∈ ℝ is a constant and 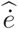 is an estimate of the derivative of the target error *e*. Since *e* = −*x* − *r* and *r* is a constant, we have 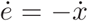. Then applying (5.14) to (3.3) and using (3.5), the closedloop system becomes:

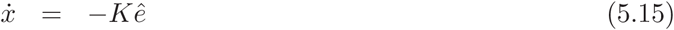

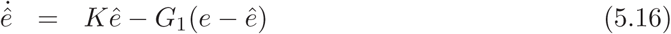

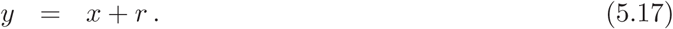

Define the *estimation error*

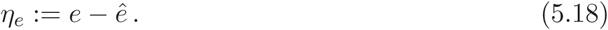

Then

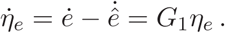

The solution of this differential equation is *η*_*e*_(*t*) = exp(*G*_1_t)*η*_*e*_(0), where *η*_*e*_(0) is the initial condition for *η*_*e*_. This gives

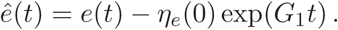

Substituting this expression into (5.15), we get

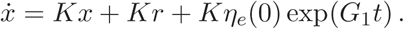

Assuming *r* is a constant, the solution of this equation is:

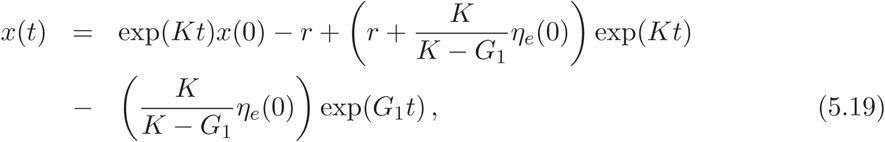

where *x*(0) is the initial condition. If we assume *r* is a constant and that the closed loop system is stable, i.e. *K* < 0 and *G*_1_ < 0, then the steadystate solution is *x*_*ss*_ ≔ lim_*t*→∞_ *x*(*t*) = −*r* and y_*ss*_ = 0. Also, the steadystate error is *e*_*ss*_ = lim_*t*→∞_ *−x(t) − r = 0*. We can see that the asymptotic behavior of this model is the one we expect. Figure 13 shows the behavior when r takes a sequence of three values: −30, 30, and 0 corresponding to a LUW experiment.

**Figure 13:**
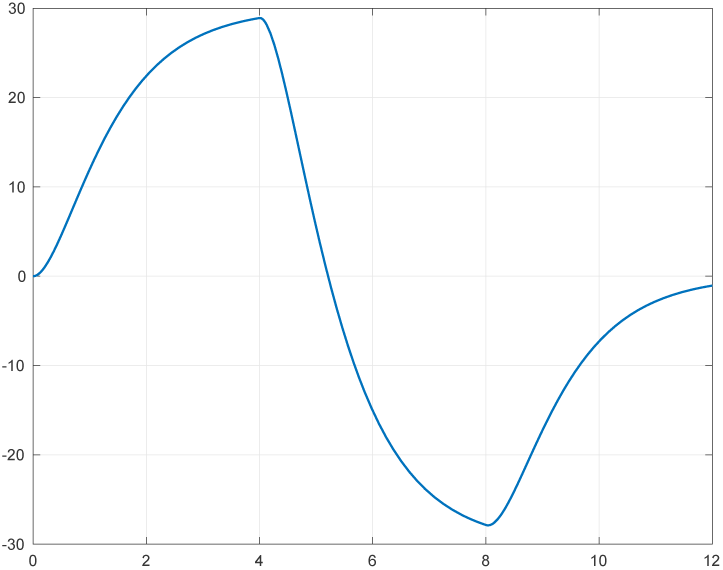
Response *x*(*t*) of (5.15)-(5.17) with *K* = −1 and *G*_1_ = −2 for an LUW experiment.

It worth noting that the response *x*(*t*) includes exponentials with two rates *K* and *G*_1_. Because of this feature, this model exhibits both savings and reduced savings. Figure 14 illustrates savings for this model. Figure 14(a) shows that in the first learning block, the time for *x*(*t*) to reach 29.75 radians starting from *x*_0_ = 0 at *t*_0_ = 0 is 5.48 seconds. In the second learning block, the time for *x*(*t*) to reach 29.74 radians starting from the same initial value is 5 seconds. Figure 14(b) shows the two responses for *r* = −30 superimposed and both starting from the initial value of 0. We can see that the rate of the response in the second block (shown in blue) is faster than the rate of the response in the first block (show in red).

**Figure 14:**
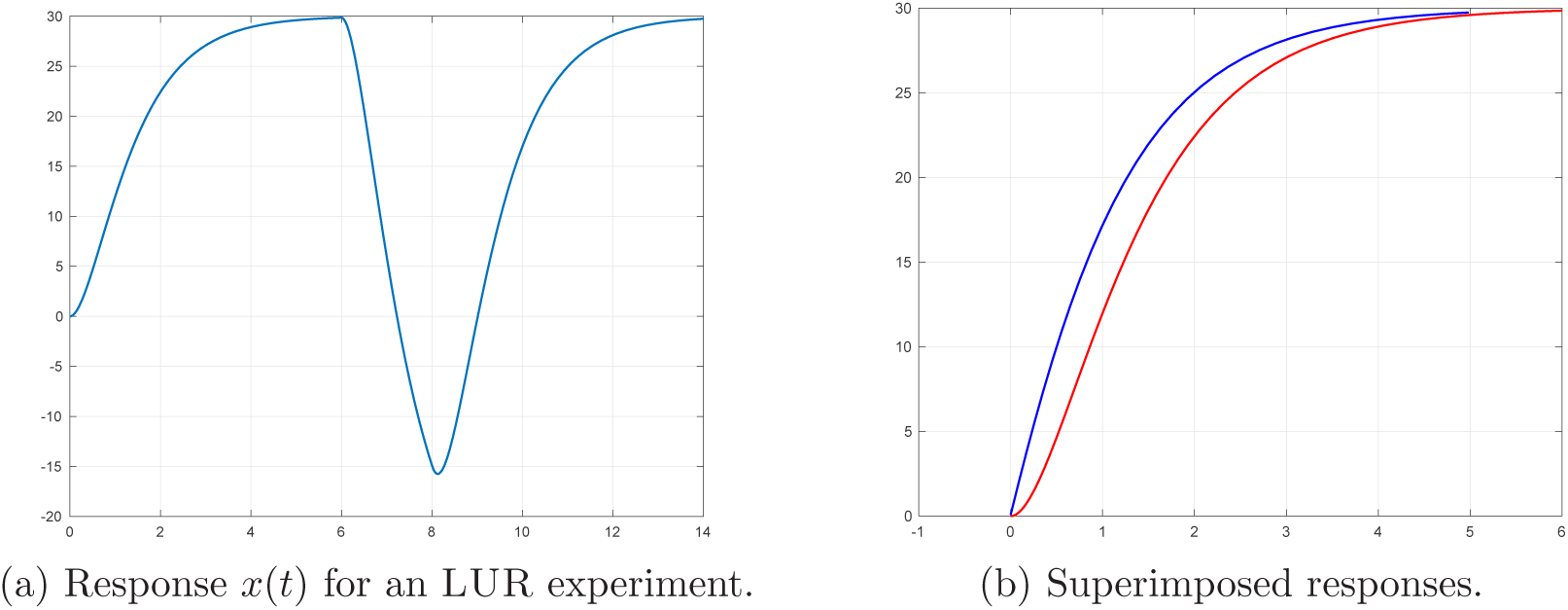
Savings experiment for (5.15)-(5.17).

The mechanism behind savings in (5.15)-(5.17) is the following. In the first learning block, the initial conditions are *x*(0) = 0 and ê(0) = 0. But in the second learning block, when *x*(*t*_1_) = 0 at time *t*_1_ = 9 seconds, we have ê(*t*_1_) = 22. This large error appears on the right hand side of (5.15) and drives 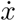 to a large value. This increase in rate ultimately leads to a decrease in rise time in the second learning block, thereby resulting in savings.

The proposed model (5.15)-(5.17) does not account for all the behaviors observed in the visuomotor rotation experiment. Here we focus on two behaviors, anterograde interference in the sense of Remark 5.2 and spontaneous recovery. Neither behavior can be generated by this model.

**Lemma 5.2.** *The stable closed-loop system* (5.15)-(5.17) *does not exhibit anterograde interference in the sense of Remark 5.2.*

*Proof.* If we compare the model (5.1)-(5.2) with (5.15)-(5.17), we observe that *n* = 2, *q* = 1, *ξ* = (x, ê), *y* = *x*, and *w* = *r*. Then we can rewrite (5.15)-(5.17) in the form of (5.1)-(5.2) to obtain the state model:

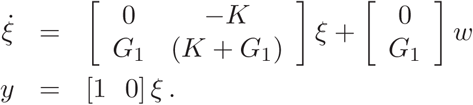

Now consider a BLU experiment. We assume without loss of generality that the learning block occurs over a time interval [0, *t*_1_) with *t*_1_ > 0 and with a constant rotation *w*(*t*) = *r* < 0 on the interval [0, *t*_1_). Unlearning occurs over the time interval [*t*_1_, ∞) with a constant rotation *w*(*t*) = −*r* > 0 on the interval [*t*_1_, ∞).

For the learning block, the effect of the baseline block is to set the initial conditions to *x*(0) = 0 and ê (0) = 0. Also *w*(*t*) = *r*, *e*(0) = −*x*(0) − *w* = −*r*, and *η*_*e*_(0) = −*r*. For the unlearning block, we assume that *t*_1_ > 0 is sufficiently large such that *x*(*t*_1_) ⋍ *x*_*ss*_ = −*r* and ê (*t*_1_) ⋍ 0. Also, *w*(*t*) = −*r*, *e*(*t*_1_) = −*x*(*t*_1_) − *w* = 2*r*, and *η*_*e*_(*t*_1_) = *e*(*t*_1_) - ê (*t*_1_) = 2*r*. Using the above information in (5.19), we obtain the solution *x*(*t*) for the learning block:

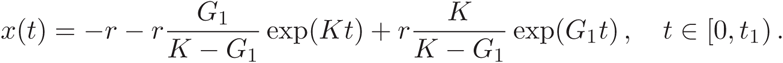

Also, the solution for *x*(*t*) for the unlearning block is:

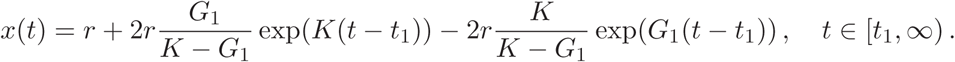

Now consider *v*(*t*) given in (5.3). We have

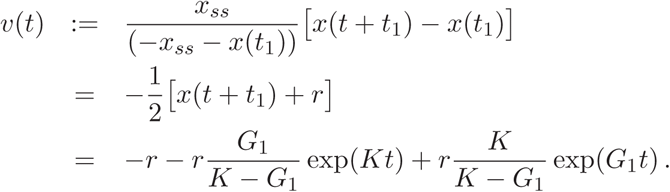

Since *x*(*t*) = *v*(*t*) for *t* ∈ [0, *t*_1_), we conclude the system (5.15)-(5.17) does not exhibit anterograde interference. □

While we have not yet discovered a rigorous proof, simulations show that for all values of *K* and *G*_1_ such that the system is stable, (5.15)-(5.17) does not exhibit anterograde interference in the sense of Definition 5.5.

**Lemma 5.3.** *The stable closed-loop system* (5.15)-(5.17) *does not exhibit spontaneous recovery.*

*Proof.* As in the proof of Lemma 5.2, we compare the model (5.1)-(5.2) with (5.15)-(5.17) and find that *n* = 2, *q* = 1, *ξ* = (*x*, ê), *y* = *x*, and *w* = *r*. Then we can rewrite (5.15)-(5.17) in the form of (5.1)-(5.2) to obtain the state model:

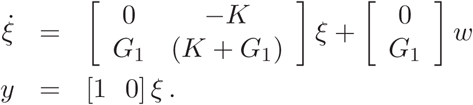

Now consider a BLUW experiment. We assume without loss of generality that the learning block occurs over a time interval [0, *t*_1_) with *t*_1_ > 0 and with a constant rotation *w*(*t*) = *r* < 0 on the interval [0, *t*_1_). Unlearning occurs over the time interval [*t*_1_, *t*_2_) with a constant rotation *w*(*t*) = −*r* > 0 on the interval [*t*_1_, *t*_2_). Finally washout occurs over the time interval [*t*_2_, ∞) with constant rotation *w*(*t*) = 0 for *t* ≥ *t*_2_.

For the learning block, the effect of the baseline block is to set the initial conditions to *x*(0) = 0 and ê(0) = 0. Also *w*(*t*) = *r*, *e*(0) = −*x*(0) − *w*(0) = −*r*, and *η*_*e*_(0) = −*r*. Using the above information in (5.19), we obtain the solution *x*(*t*) for the learning block:

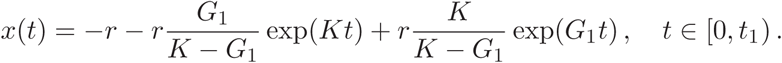

For the unlearning block, we assume that *t*_1_ > 0 is sufficiently large such that *x*(*t*_1_) ⋍ −*r* and 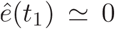. Let 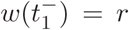 and 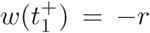 denote the value of w just before and after the time *t*_1_ when the jump in value occurs. Since *x* and ê are continuous, we have 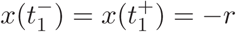 and 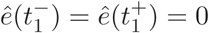. Instead, 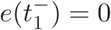 and 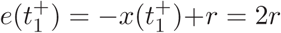. This jump in *e*(*t*) causes a jump in value of *η*_*e*_(*t*) at *t*_1_ such that 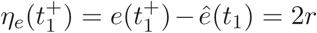. Using the above information in (5.19), we obtain the solution *x*(*t*) for the unlearning block:

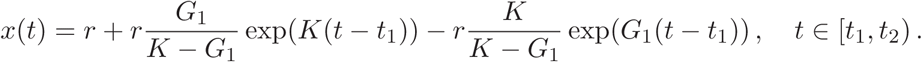

Finally, we assume that the washout block starting at *t*_2_ is such that *x*(*t*_2_) = -∊, where 0 < ∊ < −*r* and 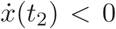 (such a time must exist since *x*(*t*) asymptotically approaches *r* < 0 in the unlearning block. That is, the washout block begins at time *t*_2_ after *x*(*t*) has crossed 0 during the unlearning block. See Figure 15 (b). Let *z*(*t*) = *x*(*t* + *t*_2_). Also let 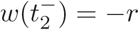 and 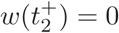 denote the value of *w* just before and after the time *t*_2_ when the jump in value occurs. Also above, we have 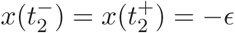 and 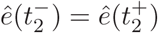. Instead, 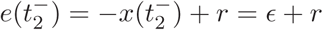 and 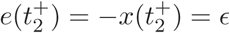. Now using (5.19), we obtain

**Figure 15:**
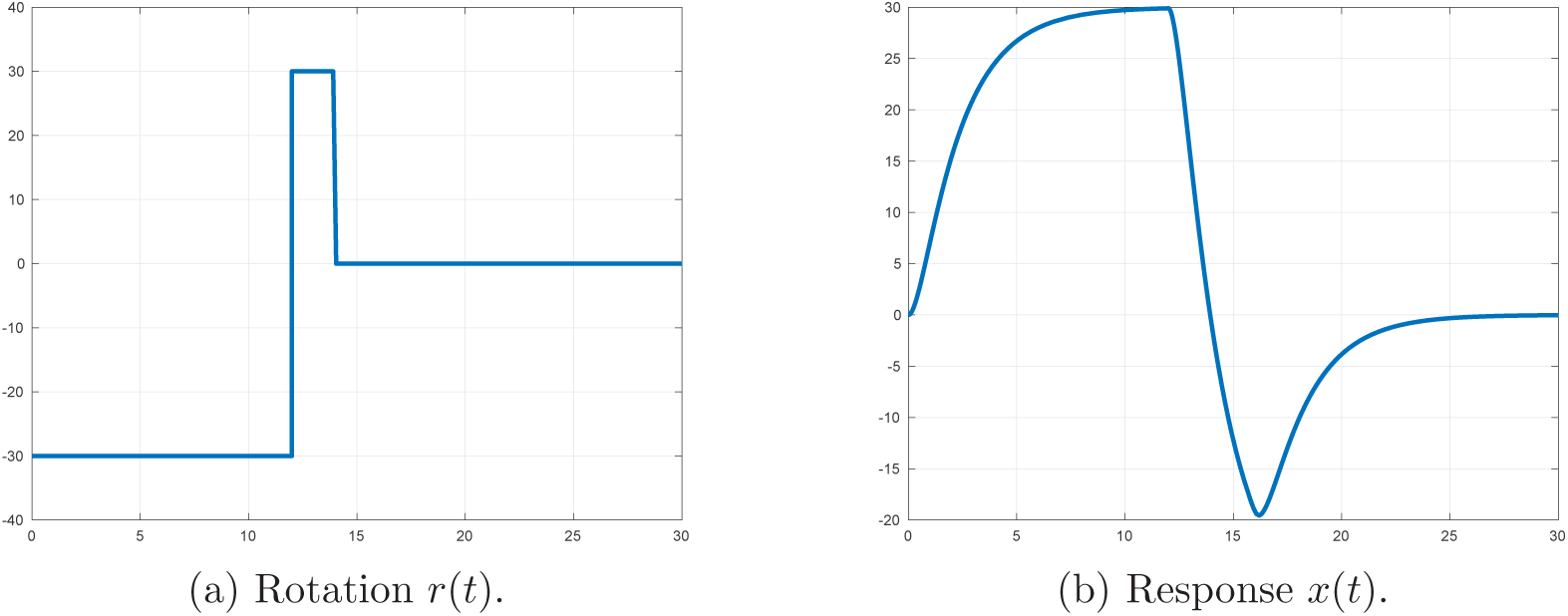
Spontaneous recovery is not exhibited by (5.15)-(5.17) in a LUW experiment.

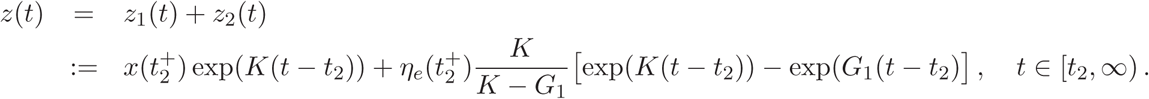

To show that the system exhibits spontaneous recovery, we must show there exists a time *t*_3_ > *t*_2_ such that *x*(*t*_3_) > 0. We argue this cannot happen by considering the individual signals *z*_1_(*t*) and *z*_2_(*t*). First, observe that since 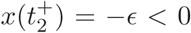 for all *t* ≥ *t*_2_. Second consider *z*_2_(*t*). First we examine

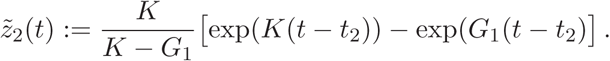

We know that *K* < 0 and *G*_1_ < 0 in order that the closed loop system is stable. If *K* < *G*_1_ < 0, then exp(*K*(*t*−*t*_2_))-exp(*G*_1_(*t*−*t*_2_) ≤ 0 for all *t* ≥ *t*_2_. Also 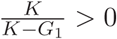. Thus, 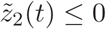 for all *t* ≥ *t*_2_. Instead if *G*_1_ < *K* < 0, then exp(*K*(*t* − *t*_2_)) - exp(*G*_1_(*t* − *t*_2_) ≥ 0 for all *t* ≥ *t*_2_. Also 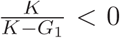. Again we find, 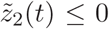 for all *t* ≥ *t*_2_. Now we have that 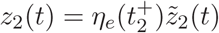. If we can show that 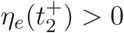, then *z*_2_(*t*) ≤ 0 for all *t* ≥ *t*_2_. Recall that 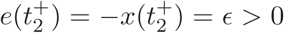. Also by assumption 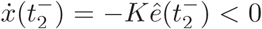, so 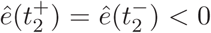 Thus, 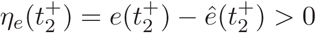, as desired.

Since *z*(*t*) = *z*_1_(*t*) + *z*_2_(*t*), we obtain *z*(*t*) ≤ 0 for all *t* ≥ *t*_2_. We conclude there is no spontaneous recovery.

From this analysis we conclude the two state model with an internal model of the error is not adequate to capture the behavior in the visuomotor experiment. Error feedback and internal models are likely components of a more sophisticated model.

### 5.5 Regulator Theory II

In this section we reapply regulator theory starting from the error feedback model (5.6)-(5.7). If we take *u* = −*Ke* + *u*^′^ as a feedback transformation with *u*^′^ a new exogenous input, apply this feedback transformation to (3.3), and rename *u*^′^ to be *u*, then we obtain a model of the form

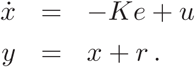

In this case, comparing (5.20)-(5.20) with the general model (A.1), we have *A* = *K*, *B* = 1, *E* = *K*, *C* = 1, *D* = 1, and *w* = *r*. The exosystem is the same as before so *S* = 0. Now we must verify conditions (N1) and (N2) to apply Theorem A.1. The pair (*A*, *B*) = (*K*, 1) is trivially controllable and therefore stabilizable. Similarly, the pair (*C*, *A*) = (1, *K*) is trivially observable and therefore detectable. Next we verify (N3). We have

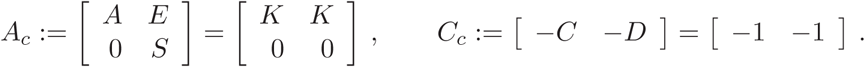

The observability matrix associated with the pair (*C*_*c*_, *A*_*c*_) is:

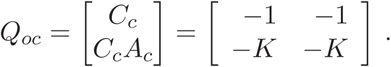

This matrix is not full rank so the pair (*C*_*c*_, *A*_*c*_) is not observable. It is also easy to check that (*C*_*c*_, *A*_*c*_) is not detectable since 0 ∈ ℂ^+^ is an unobservable eigenvalue. Instead of applying Lemma A.2 as we did above, we now proceed differently. First we solve the regulator equations (A.5)-(A.6) for the unknowns (Π, Γ). Substituting the plant and exosystem parameter values, we have

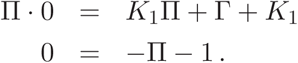

The solution of these equations is Π = −1 and Γ = 0. Under full information of plant and exosystem state, the feedback control has the form:

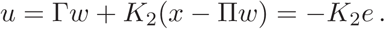

We must select *K*_2_ such that σ(*A* + *BK*_2_) ⊂ ℂ^−^. That is, *K*_1_ + *K*_2_ < 0. Since *K*_1_ < 0, we could simply choose *K*_2_ = 0. Instead we take *K*_2_ < 0. The regulator has the form of an observer as given in (A.7). We obtain

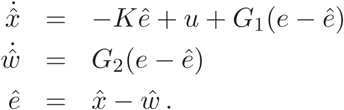

Now we observe that

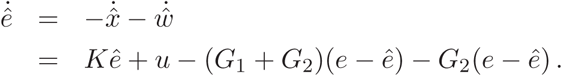

Also, the observerbased feedback is

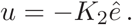

Let *K*_1_ :≔ *K* + *K*_2_ and *G* ≔ *G*_1_ + *G*_2_. The final model we obtain is

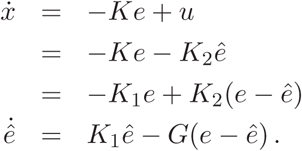

This is the model proposed in (4.3)-(4.4).

## A Regulator Theory

We review regulator theory as it pertains to the present problem. Our treatment generally follows that of [45]. We consider the LTI system

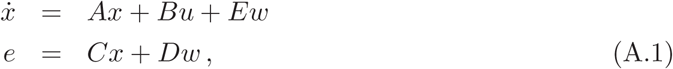

where *x* ∈ ℝ^*n*^ is the *state*, *u* ∈ ℝ^*m*^ is the *control input*, and *e* ∈ ℝ^*p*^ is the *error*. The first equation describing the system to be controlled is often called the *plant*, while the second equation describes the variable to be regulated. The terms *Ew* and *Dw* represent unmeasurable exogenous disturbance signals that enter the plant and the error. These disturbances are modeled by an *exosystem* given by

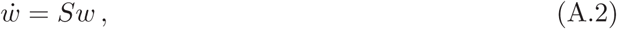

where *w* ∈ ℝ^*q*^ is the *exosystem state*. The error e also has the interpretation as the *measurement* of both the system and exosystem states. The control problem we consider is the *output regulation problem*: to design a dynamic measurement feedback of the form

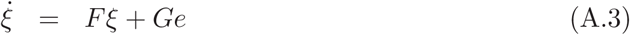

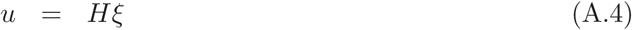

such that:

(AS) If *w*(*t*) ≡ 0, then the closed-loop system

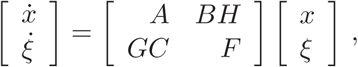

is asymptotically stable.

(R) For all initial conditions (*x*(0), *ξ*(0), *w*(0)), the closed-loop system satisfies *e*(*t)* → 0 as *t* → ∞.

A controller of the form (A.3)-(A.4) that satisfies the above objectives is called a *regulator*. Two necessary conditions to guarantee existence of a measurement feedback to stabilize the closed-loop system are:

(N1) The pair (*A*, *B*) is stabilizable.

(N2) The pair (*C*, *A*) is detectable.

### A.1 Observer-based Regulator Design

A first result on existence of a regulator is based on the idea to construct an *observer* to estimate both the system state *x* and exosystem state *w.* This construction requires an additional detectability assumption, stated below, which says that both *x* and *w* can be recovered from knowledge of the error signal *e*.

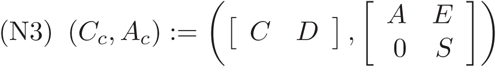 is detectable.

Using the previous assumption we get a first answer on construction of a regulator.

**Theorem A.1.** *Consider the system* (A.1) *and the exosystem* (A.2). *Suppose that* σ(S) ⊂ ℂ^+^ *and conditions (N1)(N3) hold. A regulator of the form* (A.3)(A.4) *exists if and only if there exist maps* Π : ℝ^*q*^ → ℝ^*n*^ *and* Γ : ℝ^*q*^ → ℝ^*m*^ *satisfying the* regulator equations:

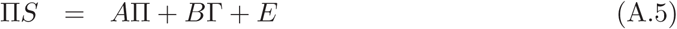

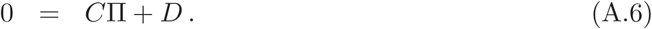

The main idea to design the regulator (A.3)-(A.4) is to construct an *observer* for the com posite state *x*_*c*_ ≔ (*x*, *w*) ∈ ℝ^*n*+*q*^. The composite system is

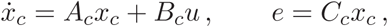

where *A*_*c*_ and *C*_*c*_ are defined above and

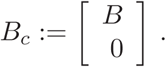

An observer for the composite system is

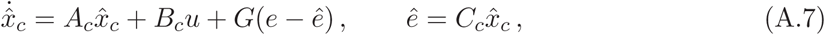

where the composite state estimate is 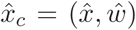. The *estimation error* is 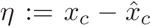, and it has dynamics

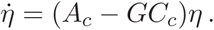

By Assumption (N3), (*C*_*c*_, *A*_*c*_) is detectable so we can choose *G* such that *A*_*c*_ − *GC*_*c*_ is Hurwitz. We will construct a feedback of a particular form:

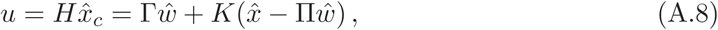

where (Π, Γ) are determined by the regulator equations. Since (*A*, *B*) is stabilizable, we choose *K* such that *A* + *BK* is Hurwitz. Now we give the proof of Theorem A.1.

*Proof.* (⟸) Suppose (Π, Γ) is a solution of (A.5)-(A.6). Consider the controller described by (A.7)-(A.8). We will show this controller is a regulator with 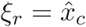. First we check the the asymptotic stability requirement (AS). Suppose *w*(*t*) ≡ 0. Then by substituting (A.8) into (A.1), the dynamics of the closed-loop system are given by

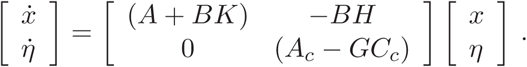

We choose *K* such that σ(*A* + *BK*) ⊂ ℂ^−^. Also we choose *G* such that σ(*A*_*c*_ − *GC*_*c*_) ⊂ ℂ^−^. Then the overall system matrix is Hurwitz. We notice that when *w*(*t*) ≡ 0, then 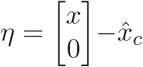, so the states (*x*, *η*) and 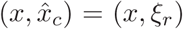 are related by a coordinate transformation. Since the system with states (*x*, *η*) (more precisely, its zero equilibrium) is asymptotically stable, so is the system with states (*x*, *ξ*_*r*_), and this proves (AS).

Second we verify the regulation requirement (R). Define *z* ≔ *x* - Π*w*. Also let 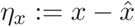 and 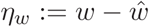, so that *η* = (*η*_*x*_, *η*_*w*_). With some algebraic manipulation we find

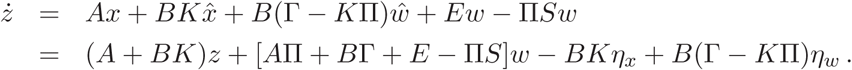

Using (A.5), we obtain

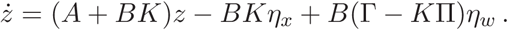

Combining with the dynamics of *η* we have the composite closed-loop dynamics

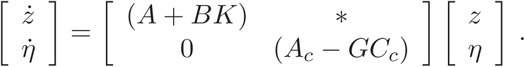

Since σ(*A* + *BK*) ∪ σ(*A*_*c*_ − *GC*_*c*_) ⊂ ℂ^−^, we have *z*(*t*) → 0. Next consider

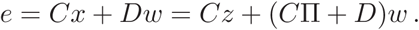

Using (A.6) we get

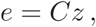

from which it follows that *e*(*t*) -→ 0, which proves (R).

(⟸) Suppose we have a regulator of the form (A.3)-(A.4). When *w*(*t*) ≡ 0, the closed-loop dynamics are

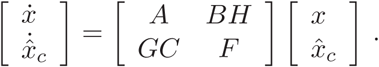

By assumption (AS), the closed-loop dynamics are asymptotically stable, so the system matrix on the right is Hurwitz. Now consider the Sylvester equation

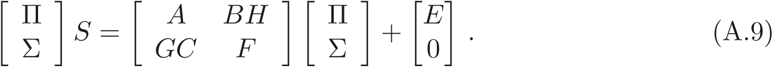

Because σ(*S*) ⊂ ℂ^+^ and the closedloop system matrix is Hurwitz, their spectra are disjoint. Then by Sylvester’s theorem [20] there exists a unique solution for (Π, σ). Write the matrix equation (A.9) as two equations:

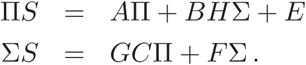

In particular, if we set Γ ≔ *H*σ, then we obtain (A.5). Next let *z*_1_ = *x*−Π*w* and *z*_2_ = *ξ*−σ*w*. Then using (A.9) we obtain

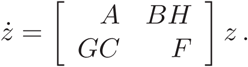

Since this system matrix is Hurwitz based on the observation above, *z*(*t*) → 0. Now consider

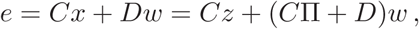

as above. By assumption *e(t*) → 0. Also, *z*(*t*) → 0. Therefore, (*C*Π + *D*)*w*(*t*) -→ 0 for all initial conditions *w*(0). However, eig(*S*) ⊂ ℂ^+^, so there exists an initial condition of the exosystem *w*(0) such that the corresponding solution satisfies 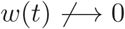. This implies *C*Π + *D* = 0, which proves (A.6). □

The previous result was based on a detectability assumption (N3) that allowed an observer to be constructed to estimate *x* and *w*. This assumption is quite restrictive, and indeed will not be met in our application of regulator theory. On the other hand, (N1) and (N2) are necessary conditions. Now we give a procedure to solve the problem when (N1) and (N2) hold, but (N3) does not. In this case, the system is detectable by (N2) but the combined plant and exosystem is not detectable. The idea is to perform a coordinate transformation and then trim off the part of the exosystem model that is redundant with the system and causes the failure of detectability.

**Lemma A.2.** *Consider the system* (A.1) *and the exosystem* (A.2). *Suppose that conditions (N1) and (N2) hold but not (N3). Consider the composite system*

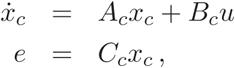

*where x*_*c*_ = (*x*, *w*), *and A*_*c*_, *B*_*c*_, *and C*_*c*_ *are defined above. There exists a coordinate trans-formation* 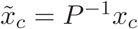 *such that in new coordinates the composite system is:*

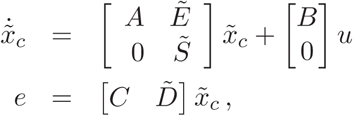

*where* 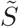, 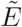, *and* 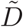 *have a partitioned structure*

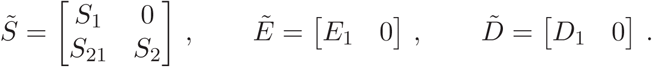

*Moreover, the pair* 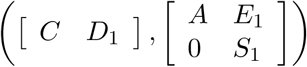 *is detectable.*

*Proof.* It is wellknown that if the pair (*C*_*c*_, *A*_*c*_) is not detectable, then there exists a co-ordinate transformation 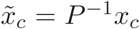 such that in new coordinates the transformed system is:

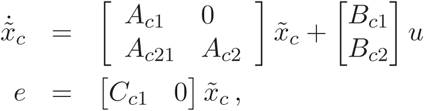

where (*C*_*c*1_, *A*_*c*1_) is detectable. Since the pair (*C*, *A*) is detectable, this transformation can be chosen such that

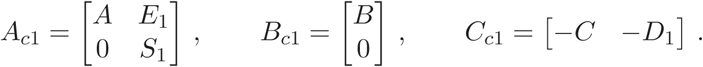

Lemma A.2 gives a procedure to resolve the problem of (N3) failing for the original system. First apply the coordinate transformation 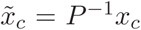, where 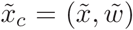. If we partition 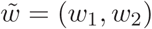 as in the proof, then the reduced system is:

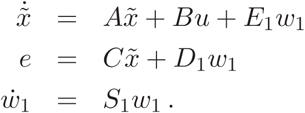

We can see that the original plant is retained, but the exosystem has been trimmed. The resulting system satisfies (N3) so we can apply Theorem A.1 to design a regulator.

*Remark* A.1. It is possible that in applying the reduction process of Lemma A.2, the com-ponent *w*_1_ of the exosystem state is vacuous. In this case the output regulation problem reduces to a problem of stabilizing the plant of the reduced system.

### A.2 Structural Stability

The previous results were based on the idea that the system matrices {*A, B, C, D, E*} are all exactly known. In practice, these parameters are only approximately known, yet we would still like to construct a regulator to solve the output regulation problem. These leads us to the problem of a *structurally stable synthesis*. We begin by viewing the set of matrices {*A, B, C, D, E*} as elements of a space of parameters

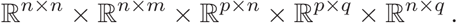

Uncertainty in these parameters is modeled as the set {*A, B, C, D, E*} varying over a neigh-borhood 𝒫_0_ of the nominal parameter values {*A*_0_, *B*_0_, *C*_0_, *D*_0_, *E*_0_}.

**Definition A.1.** *A* controller of the form (A.3)-(A.4) is called a *robust regulator* at {*A*_0_, *B*_0_, *C*_0_, *D*_0_, *E*_0_}if

(i) It solves the output regulation problem for the nominal parameter values {*A*_0_, *B*_0_, *C*_0_, *D*_0_, *E*_0_}.

(ii) It solves the output regulation problem for each perturbed set of parameter values {*A, B, C, D, E*} so long as the system matrix 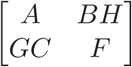 is Hurwitz

We consider a nomimal plant and an exosystem

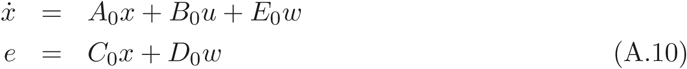

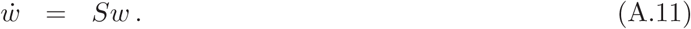

It is assumed that *S* is a known, constant matrix, unlike the plant parameters {*A*_0_, *B*_0_, *C*_0_, *D*_0_, *E*_0_} that are not known exactly. The main result on existence of a robust regulator in the SISO case that we study is the following.

**Theorem A.3.** *Consider the nominal system* (A.10) *and the exosystem* (A.11) *with* m = *p* = *q* = 1. *Suppose that* σ(S) ⊂ ℂ^+^. *There exists a robust regulator if and only if the pair* (*A*_0_, *B*_0_) *is stabilizable, the pair* (*C*_0_, *A*_0_) *is detectable, and the matrix*

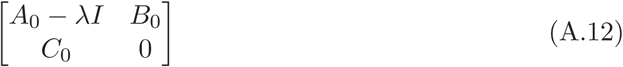

*is nonsingular for each* λ ∈ σ(*S*).

In light of the previous result and to simplify some steps of the design, we make the following assumptions.

**Assumption A.1.** Consider the nominal system (A.10) and the exosystem (A.11).

(A0) *m* = *p* = *q* = 1.

(A1) The pair (*A*_0_, *B*_0_) is stabilizable.

(A2) The pair (*C*_0_, *A*_0_) is detectable.

(A3) The pair (D_0_, *S*) is detectable.

(A4) The exosystem satisfies σ(S) ⊂ ℂ^+^.

(A5) σ(*S*) ∩ σ(*A*_0_) = ∅.

The proof of sufficiency of Theorem A.3 is by construction. We begin with an auxiliary system to which we wish to apply Theorem A.1. The *auxiliary system* is

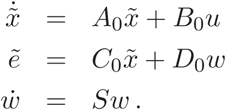

The next result shows that the auxiliary system satisfies condition (N3).

**Lemma A.4.** *Consider the nominal system* (A.10) *and the exosystem* (A.11). *Suppposethat Assumption A.1 holds. Then the pair* 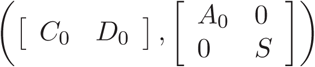 *is detectable.*

*Proof.* This result comes from the fact that (*C*_0_, *A*_0_) is detectable, (*D*_0_, *S*) is detectable, and σ(*S*) ∩ σ(*A*_0_) = ∅.

Since the auxiliary system (A.13)-(A.13) satisfies (N3) as well as (N1)-(N2) by assumptions (A1) and (A2), we can use Theorem A.1 to design a regulator for the auxiliary system. The regulator has the general form

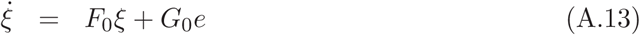

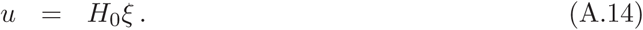

In more detail, let 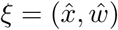. Then

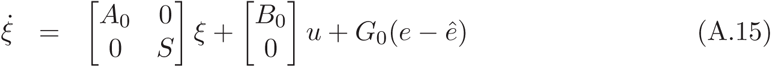

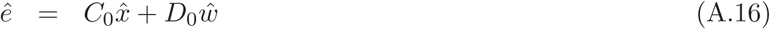

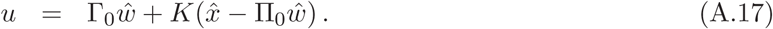

Here we choose *K* such that σ(*A*_0_ + *B*_0_*K*) ⊂ ℂ^−^. Also, we assume (Π_0_, Γ_0_) is a solution of

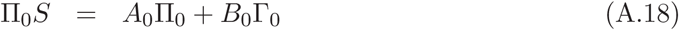

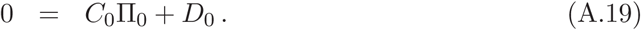

When *w*(*t*) ≡ 0, the closed-loop system is

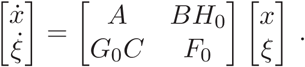

In order for the regulator to satisfy condition (*S*), we assume

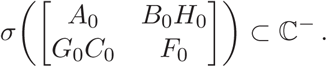

Now consider a perturbed plant

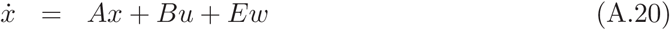

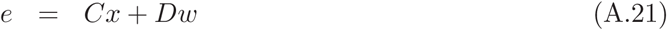

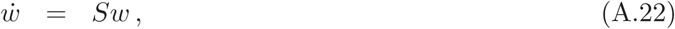

where *x* ∈ ℝ^*n*^>, *u* ∈ ℝ^*m*^, *e* ∈ ℝ^*p*^, and *w* ∈ ℝ^*q*^. We use the same controller given in (A.15) (A.17). We assume that perturbation on the plant parameters is restricted so that when *w*(*t*) ≡ 0, the closed loop system matrix with the perturbed plant satisfies

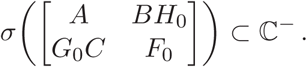

This assumption will address the stability requirement (S) of a regulator for the perturbed plant. It then remains to show that the regulation requirement (R) is satisfied. The next two results address this requirement.

**Lemma A.5.** *Consider the nomimal system* (A.10), *the perturbed system* (A.20), *and the exosystem* (A.22). *Suppose that Assumption A.1 holds. Let* (A.13)-(A.14) *be a regulator for the nomimal system* (A.10). *Suppose 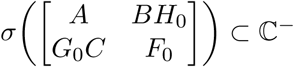. Then there exist* (Π, σ) *such that*

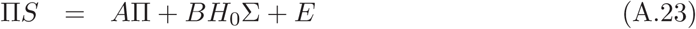

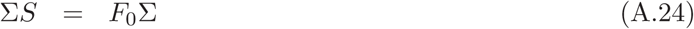

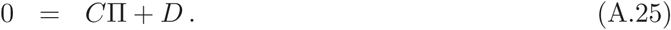

*Proof.* Since 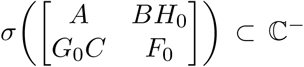 and σ(S) ⊂ ℂ^+^, by Sylvester’s Theorem [20] the

Sylvester equation

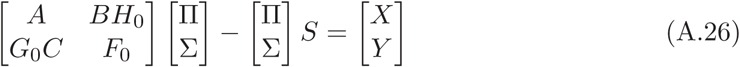

is solvable in (Π, σ) for each (*X*, *Y*). We choose *X* = −*E* and *Y* = −*G*_0_*D*. Then the previous equation gives

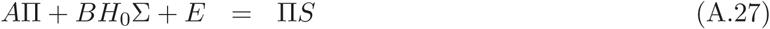

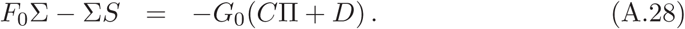

The first equation (A.27) gives (A.23). To understand (A.28), we define two linear maps ℒ _1_ : ℝ^2^ → ℝ^2^ and ℒ _2_ : ℝ^1^ → ℝ^2^ given by

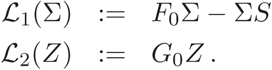

Then (A.28) says that

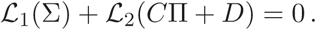

If we can show that Im ℒ _1_ ∩ Im ℒ _2_ = {0} and Ker ℒ _2_ = {0}, then (A.24)-(A.25) follow. First, using (A.15)-(A.17), we observe that, in more detail, (A.24) is equivalent to the equation

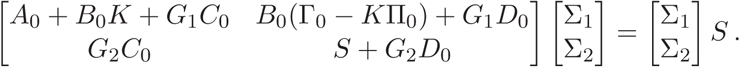

That is,

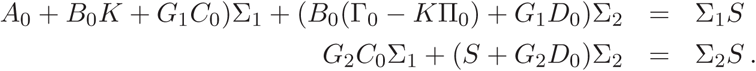

Now we claim that 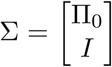, or equivalently, is a solution of *F*^0^ σ = σ*S*. If we substitute σ_1_ = Π_0_ and σ_2_ = *I* in the left hand side of the equation above and we use the fact that (Π_0_, Γ_0_) are solutions of (A.18)-(A.19), we get

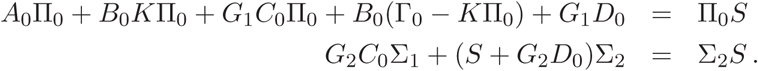

This shows that 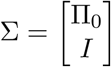 is a solution of (A.24). This implies that dim(Ker ℒ _1_) ≥ 1. But we know that ℒ _1_ : ℝ^2^ → ℝ^2^ so

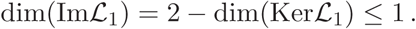

Also since ℒ _2_ : ℝ^1^ → ℝ^2^, dim(Im ℒ _2_) ≤ 1.

We already know from our discussion of (A.26) that for all *Y*, there exist (Π, σ) such that

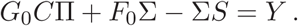

Equivalently, ℒ_1_(σ)+ ℒ_2_(CΠ) = *Y*. Since *Y* is arbitrary, we deduce that Im ℒ_1_ +Im ℒ_2_ = ℝ^2^. Combined with the statements above, we find dim(Im ℒ_1_) = 1, dim(Im ℒ_2_) = 1, Im ℒ_1_ ∩ Im ℒ_2_ = {0} and dim(Ker ℒ_2_)0, as desired. □

**Lemma A.6.** *Consider the system* (A.20)-(A.22) *and the controller* (A.13)-(A.14). *Sup pose that* 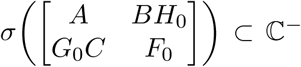 *and* σ(S) ⊂ ℂ^+^. *Let* (Π, σ) *be a solution of* (A.23)- (A.25). *Then for all* (*x*(0), *ξ*(0), *w*(0)), *e*(*t*) → 0 *as t* → ∞.

*Proof.* a Let (Π, σ) be the solution of (A.23)-(A.25). Define *z* ≔ *x* - Π*w*. Then

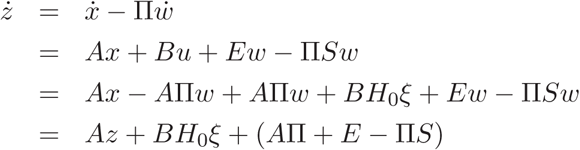

Let 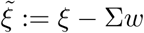. Then we have

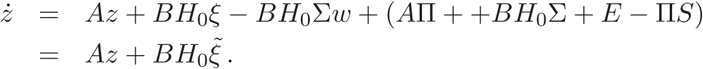

Next consider

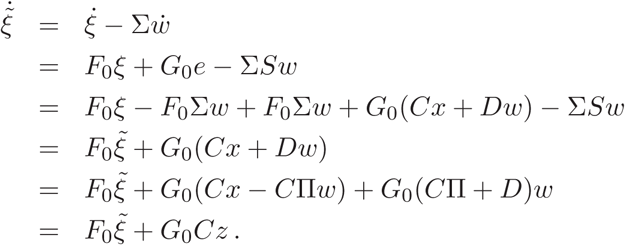

The overall system is

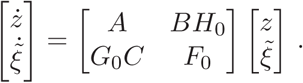

By assumption σ() ⊂ ℂ^−^. Therefore *z*(*t*) → 0. However, we know that

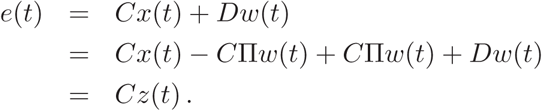

We conclude that *e*(*t*) → 0, as desired. □

## References

[1] J. Albus. A theory of cerebellar function. Mathematical Biosciences. Vol. 10, No 1, pp. 25–61, February 1971.

[2] A. Ahmed, D. Wolpert, and J. Flanagan. Flexible representations of dynamics are used in object manipulation. Curr Biology. Vol. 18, No. 10, pp. 763768, 2008.

[3] B. Benson, J. Anguera, R. Seidler. A spatial explicit strategy reduces error but inter feres with sensorimotor adaptation. J Neurophysiology. Vol. 105, pp. 28432851, 2011.

[4] M. Berniker and K. Kording. Estimating the sources of motor errors for adaptation and generalization. Nature Neuroscience. No. 11, pp. 14541461, 2008.

[5] M. Berniker and K. Kording. Estimating the relevance of world disturbances to explain savings, interference, and longterm adaptation effects. PLoS Computational Biology. Vol. 7, issue 10, October 2011.

[6] K. Bond and J. Taylor. Flexible explicit but rigid implicit learning in a visuomotor adaptation task. J. Neurophysiology. No. 113, pp. 38363849, 2015.

[7] V. Braitenberg. Is the cerebellar cortex a biological clock in the millisecond range? Progress in Brain Research. Vol. 25, 334–346, 1967.

[8] P. Butcher and J. Taylor. Decomposition of a sensory prediction error signal for visuo-motor adaptation. J. Experimental Psychology: Human Perception and Performance. May 2017.

[9] S. Cheng and P. Sabes. Modeling sensorimotor learning with linear dynamical systems. Neural Computation. Vol. 18, pp. 760793, 1996.

[10] M. Conditt, F. Gandolfo, F. MussaIvaldi. The motor system does not learn the dy-namics of the arm by rote memorization of past experience. J Neurophysiology. Vol. 78, No. 1, pp. 55460, 1997.

[11] H. Cunningham. Aiming error under transformed spatial mappings suggests a structure for visual-motor maps. J. Experimental Psychology. Vol. 15, No. 3, pp. 493506, 1989.

[12] E.J. Davison and A. Goldenberg. Robust control of a general servomechanism problem: the servo compensator. Automatica. Vol. 11, No. 5, pp. 461–471, 1975.

[13] J. Eccles, M. Ito, and J. Szentagothai. The Cerebellum as a Neuronal Machine. SpringerVerlag, 1967.

[14] V. Ethier, D. Zee, and R. Shadmehr. Spontaneous recovery of motor memory during saccade adaptation. J. Neurophysiology. Vol. 99, May 2008.

[15] B.A. Francis. The linear multivariable regulator problem. SIAM J. Control and Opti-mization. Vol. 15, No. 3, pp. 486–505, May 1977.

[16] B.A. Francis and W.M. Wonham. The internal model principle for linear multivariable regulators. Applied Mathematics and Optimization. Vol. 2, No. 2, 1975.

[17] B.A. Francis and W.M. Wonham. The internal model principle of control theory. Au-tomatica. Vol. 12, pp. 457–465, 1976.

[18] D. Franklin, A. Reichenbach, S. Franklin, and J. Diedrichsen. Temporal evolution of spatial computations for visuomotor control. J. Neuroscience. Vol. 36, No. 8, pp. 2329– 2341, February 2016.

[19] J. Galea, A. Vazquez, N. Pasricha, J. de Xivry, P. Celnik. Dissociating the roles of the cerebellum and motor cortex during adaptive learning: the motor cortex retains what the cerebellum learns. Cerebral Cortex. No. 21, pp. 1761–1770, 2011.

[20] F.R. Gantmacher. The Theory of Matrices. Chelsea Publishing. 1959.

[21] A. Haith and J. Krakauer. Modelbased and model-free mechanisms of human motor learning. Adv Exp Med Biology. No. 782, pp. 121, 2013.

[22] M. Haruno, D. Wolpert, and M. Kawato. MOSAIC model for sensorimotor learning and control. Neural Computation. Vol. 13, pp. 2201–2220, 2001.

[23] R. Held and N. Gottlieb. Technique for studying adaptation to disarranged hand-eye coordination. Percept. Mot. Skills. No. 8, pp. 8386, 1958.

[24] D. Herzfeld, P. Vaswani, M. Marko, and R. Shadmehr. A memory of errors in sensori-motor learning. Science. No. 345, pp. 1349–1353, 2014.

[25] V. Huang, A. Haith, P. Mazzoni, and J. Krakauer. Rethinking motor learning and savings in adaptation paradigms: modelfree memory for successful actions combines with internal models. Neuron. Vol. 70, issue. 4, pp. 787–801, May 2011.

[26] M. Kawato. Internal models for motor control and trajectory planning. Current Opinion in Neurobiology. Vol. 9, No. 6, pp. 718727, December 1999.

[27] H. Kim, J.R. Morehead, D. Parvin, R. Moazzezi, and R. Ivry. Invariant errors reveal limitations in motor correction rather than constraints on error sensitivity. bioRxiv 189597, September 2017.

[28] Y. Kojima, Y. Iwamoto, and K. Yoshida. Memory of learning facilitates saccadic adap-tation in the monkey. The Journal of Neuroscience. Vol. 24, No. 34, pp. 7531–7539, August 2004.

[29] K. Kording, J. Tenenbaum, and R. Shadmehr. The dynamics of memory as a conse-quence of optimal adaptation to a changing body. Nature Neuroscience. Vol. 10, No. 6, June 2007.

[30] J. Krakauer. Motor learning and consolidation: the case of visuomotor rotation. Ad-vances in Experimental and Medical Biology. No. 629, pp. 405421, 2009.

[31] J. Krakauer, C. Ghez, and M. Ghilardi. Adaptation to visuomotor transformations: consolidation, interference, and forgetting. J. Neuroscience. Vol. 25, No. 2, pp. |p473478, January 2005.

[32] J. Krakauer, M. Ghilardi, and C. Ghez. Independent learning of internal models for kinematic and dynamic control of reaching. Nature Neuroscience. Vol. 2, No. 11, Novem-ber 1999.

[33] J. Krakauer and P. Mazzoni. Human sensorimotor learning: adaptation, skill, and beyond. Current Opinion in Neurobiology. Vol. 21, pp. 19, 2011.

[34] J. Lee, N. Schweighofer. Dual adaptation supports a parallel architecture of motor memory. J Neuroscience. Vol. 29, pp. p. 1039610404, 2009.

[35] L. Leow, A. Rugy, W. Marinovic, S. Riek, and T. Carroll. Savings for Visuomotor adaptation require prior history of error, not prior repetition of successful actions. J. Neurophysiology.No. 116, pp. 1603–1614, 2016.

[36] D. Marr. A theory of the cerebellar cortex. Journal of Physiology. No. 202, pp. 437–470, 1969.

[37] T. Martin, J. Keating, H. Goodkin, A. Bastian, W. Thach. Throwing while looking through prisms. Brain. No. 119, pp. 1183–1198, 1996.

[38] F. Mawase, L. Shmuelof, S. BarHaim, A. Karniel. Savings in locomotor adaptation explained by changes in learning parameters following initial adaptation. J. Neurophys-iology. No. 111, pp. 14441454, January 2014.

[39] P. Mazzoni J. Krakauer. An implicit plan overrides an explicit strategy during visuo-motor adaptation. J NeuroscienceVol. 26, pp. 36423645, 2006.

[40] S. McDougle, K. Bond, and J. Taylor. Explicit and implicit processes constitute the fast and slow processes of sensorimotor learning. Journal of Neuroscience, Vol. 35, pp. 95689579, 2015.

[41] R.C. Miall and D.M. Wolpert. Forward models for physiological motor control. Neural Networks. Vol. 9, No. 8, pp. 1265–1279, 1996.

[42] J.R. Morehead, J. Taylor, D. Parvin, and R. Ivry. Characteristics of implicit sen-sorimotor adaptation revealed by taskirrelevant clamped feedback. J. of Cognitive Neuroscience. Vol. 29, No. 6, pp. 1061–1074, June 2017.

[43] R. Morehead, J. Taylor, D. Parvin, E. Marrone, and R. Ivry. Implicit adaptation via visual error clamp. In Translational and Computational Motor Control. 2014.

[44] R.A. Rescorla. Spontaneous recovery. Learning and Memory. Vol. 11, pp. 501–509, 2004.

[45] A. Saberi, A. Stoorvogel, and P. Sannuti. Control of Linear Systems with Regulation and Input Constraints. Springer, 2000.

[46] P. Sabes. The planning and control of reaching movements. Curren Opinion in Neuro-biology. No. 10, pp. 740746, 2000.

[47] R. Shadmehr, M. Smith, and J. Krakauer. Error correction, sensory prediction, and adaptation in motor control. Annual Rev. Neuroscience. Vol. 33, pp. 89108, 2010.

[48] R. Shadmehr and S. Wise. Computational Neurobiology of Reaching and Pointing: A Foundation for Motor Learning. MIT Press, 2005.

[49] G. Sing and M. Smith. Reduction in learning rates associated with anterograde in-terference results from interactions between different timescales in motor adaptation. PLoS Computational Biology. Vol. 6, Issue 8, August 2010.

[50] M. Smith, A. Ghazizadeh, and R. Shadmehr. Interacting adaptive processes with differ-ent timescales underlie shortterm motor learning. PLoS Computational Biology. Vol. 4, Issue 6, June 2006.

[51] R. Sperry. Effect of 180 degree rotation of the retinal field on visuomotor coordination. J. Experimental Zoology, No. 92, pp. 263–279, 1943.

[52] R. Sperry. Neural basis of the spontaneous optokinetic response produced by visual inversion. pp. 482 – 489.

[53] N Stollhoff, R. Menzel, and D. Eisenhardt. Spontaneous recovery from extinction depends on the reconsolidation of the acquisition memory in an appetitive learning paradigm in the honeybee. J. Neuroscience. Vol. 25, pp. 44854492, 2005.

[54] K. Takiyama, M. Hirashima, and D. Nozaki. Prospective errors determine motor learn-ing. Nature Communications. Vol. 6, January 2015.

[55] J. Taylor and R. Ivry. Flexible cognitive strategies during motor learning. PLoS Com-putational Biology. Vol. 7, Issue 3, March 2011.

[56] J. Taylor, J. Krakauer, R. Ivry. Explicit and implicit contributions to learning in a sensorimotor adaptation task. J Neuroscience. Vol. 34, pp. 3023–3032, 2014.

[57] P. Vaswani, L. Shmuelof, A. Haith, R. Deknicki, V. Huang, P. Mazzoni, R. Shadmehr, and J. Krakauer. Persistent residual errors in in motor adaptation tasks: reversion to baseline and exploratory escape. J. Neuroscience. Vol. 35, No. 17, pp. 6969–6977, April 2015.

[58] K. Wei and K. Kording. Relevance of error: what drives motor adaptation? J. Neuro-physiology. Vol. 101, pp. 655664, 2009.

[59] R. Welch, C. Choe, and D. Heinrich. Evidence for a threecomponent model of prism adaptation. J. Exp. Psychol. No. 103, issue 4, pp. 700705. 1974.

[60] V. Wigmore, C. Tong, and J.R. Flanagan. Visuomotor rotations of varying size and direction compete for a single internal model in motor working memory. J. of Experi-mental Psychology: Human Perception and Performance. Vol. 29, No. 2, pp. 447–457, 2002.

[61] D. Wolpert, Z. Ghahramani, and M. Jordan. An internal model for sensorimotor inte-gration. Science, Vol. 269, No. 5232, pp. 18801882, September 1995.

[62] D. Wolpert and M. Kawato. Multiple paired forward and inverse models for motor control. Neural Networks. Vol. 11, pp. 1317–1329, 1998.

[63] D. Wolpert, R.C. Miall, and M. Kawato. Internal models in the cerebellum. Trends in Cognitive Sciences. Vol. 2, No. 9, pp. 338–347, September 1998.

[64] W.M. Wonham. Linear Multivariable Control: A Geometric Approach. 3rd Ed. Springer-Verlag, 1979.

[65] E. Zarahn, G. Weston, J. Liang, P. Mazzoni, and J. Krakauer. Explaining savings for visuomotor adaptation: linear timeinvariant statespace models are not sufficient. J. Neurophysiology. Vol. 100, pp. 2537–2548, November 2008.

